# Alteration of pro-carcinogenic gut microbiota is associated with clear cell renal cell carcinoma tumorigenesis

**DOI:** 10.1101/2022.12.07.519551

**Authors:** Bo-Yu Yang, Fang-Zhou Zhao, Xuan-Hao Li, Mei-Shan Zhao, Jing-Cheng Lv, Ming-Jun Shi, Jun Li, Zhi-Yuan Zhou, Jing-Jing Wang, Jian Song

**Author notes:** Correspondence to: Dr. Zhi-Yuan Zhou, Dr. Jing-Jing Wang and Dr. Jian Song: **Zhi-Yuan Zhou**, GongLi Hospital of Shanghai Pudong New Area, No. 219, Miaopu Road, Shanghai 200135; **Jing-Jing Wang**, Shanghai Key Laboratory for Pancreatic Diseases, Institute of Translational Medicine, Shanghai General Hospital, Shanghai Jiao Tong University School of Medicine, 650 Xin Songjiang Road, Shanghai 201620, China; **Jian Song**, Department of Urology, Beijing Friendship Hospital, Capital Medical University, No. 95, Yongan Road, Xicheng District, Beijing100050, People’s Republic of China. These authors contributed equally to this work (Bo-Yu Yang, Fang-Zhou Zhao, Xuan-Hao Li).

## Abstract

Increasing evidence suggests that gut microbiota is involved in the occurrence and progression of urinary system diseases such as clear cell renal cell carcinoma (ccRCC). However, the mechanism of how alteration of gut metagenome promotes ccRCC remains unclear. Here we aim to elucidate the association of specific gut bacteria and their metabolites with ccRCC. In a pilot case-control study among 30 ccRCC patients and 30 healthy controls, 16S ribosomal RNA (rRNA) gene sequencing were analyzed from fecal samples collected prior to surgery or hospitalization. Alpha diversity and beta diversity analysis of the gut microbiota were performed, and differential taxa were identified by multivariate statistics. Meanwhile, serum metabolism was measured by UHPLC-MS, and differential genes were identified based on the *TCGA* database. Random Forests revealed the relative abundances of 20 species differed significantly between the RCC group and the Control group, among which 9 species, such as *Desulfovibrionaceae,* were enriched in the RCC group, and 11 species, such as four kinds of *Lactobacillus,* were less abundant. Concomitantly, serum level of taurine, which was considered to be consumed by *Desulfovibrionaceae* and released by *Lactobacillus*, has decreased in the RCC group. In addition, macrophage-related genes such as *Gabbr1* was upregulated in ccRCC patients from our results.

**IMPORTANCE:** To our knowledge, few studies investigate the correlation of gut microbiota and ccRCC tumorigenesis. Overall, our sequencing data suggest that changes in the composition of specific gut microbiota, especially *Lactobacillus* and *Desulfovibrionaceae,* may be involved in ccRCC. Numerous serum metabolites, for example, taurine, which were modified in concert with dysregulation of gut microbiota, were associated with metabolic status during ccRCC development. Furthermore, through comparative analysis of clinical indicators, we found that gut dysbiosis could potentially reshape systemic inflammation, which participated in ccRCC tumorigenesis and we performed bioinformatics analysis to draw this conclusion. In Summary, it could be concluded from our study that the reduction of protective bacteria *Lactobacillus*, proliferation of sulfide-degrading bacteria *Desulfovibrionaceae*, reduction of taurine, and enrichment of macrophage related genes might be the risk predictors of ccRCC.

---

The human gut microbiota is a complex micro-ecosystem that is closely related to human health and disease. Various types of gut microbes interact with each other and jointly maintain the normal structure of the gut by forming a bacterial barrier. They protect human body from microbial infection, participate in digestion, absorption and metabolism of nutrients, as well as regulate the gut immune response (1). Many studies have shown that the occurrence of renal related diseases is accompanied by changes in the gut microbiota characteristics, of which the metabolic status is the key factor affecting the progression of the disease.

Increasing evidence suggests that the interaction between gut microbiota and metabolic status within host has the potential to promote or influence chronic kidney disease (CKD). Recent studies proved a dual-directional regulatory relationship between gut microbiota and the host with CKD. Abnormal renal function may result from long-term effects of excess uremic toxins such as indole sulfate (IS), p-cresol sulfate (PS) and trimethylamine-N-oxide (TMAO) which are produced due to altered composition of gut microbiota. It is well known that abnormal renal function, especially CKD, are associated with oxidative stress, endotoxemia, inflammation, and a higher prevalence of cardiovascular disease, where gut dysbiosis is one of the main causes of these symptom. Additionally, gut microbiota can reduce oxidative stress-induced kidney damage by secreting short-chain fatty acids (SCFA) (2). The gut dysbiosis and subsequent leakage of pro-inflammatory products such as interleukin-6 (IL-6) and monocyte chemoattractant protein-1 (MCP-1) may result in chronic inflammatory state, which contribute to CKD (3, 4). 16S rRNA gene sequencing was performed to the feces of 13 patients with multiple kidney stones and 13 healthy controls. Results showed that there were significant differences in β-diversity and the relative abundance of 20 bacterial genera, for example *Phascolarctobacterium*, *Parasutterella*, *Ruminiclostridium*, *Erysipelatoclostridium*, *Fusicatenibacter* and *Dorea.* These bacterial genera were associated with blood concentrations of trace elements such as potassium, sodium, calcium and chlorine (5). Thus, there may exist a gut-kidney axis mediated by metabolism and inflammation whereby alterations in gut microbiota composition can affect the state of renal physiology and pathology.

The etiology of ccRCC, the most common malignant disease of the kidney, is still unclear. Studies reveal that metabolism of tryptophan, arginine and glutamine participates in the progression of ccRCC, and targeting the metabolic reprogramming of tricarboxylic acid cycle (TCA) that affect neoplastic biosynthesis has been gradually evolving as the new therapeutic strategies (6). Downregulation of conventional metabolites genes involved in lipid and amino acid biosynthesis, particularly succinate, an intermediate in the TCA, is associated with ccRCC and poor prognosis (7). Meanwhile, current studies have shown that inflammatory factors such as IL-6, IL-10, IL-1β and estrogen play an important role in the occurrence of ccRCC (8–11). Increased expression of inflammatory chemokine IL-8 and its receptor CXCR1 has been demonstrated to be associated with ccRCC tumorigenesis and decreased overall survival via Epithelial-mesenchymal transition (EMT) pathway (12).

Differences in gut microbiota compositions and functions have been proven to influence an individual’s immune system, for example the expression of inflammatory factors and the immune system’s response to pathogens (13). Gut microbiota have been verified to systematically regulate the progression of non-gastrointestinal tumors such as melanoma, lung cancer, and breast cancer through the metabolism- and inflammation-related pathways (14–16). These previous studies have provided promising insights into the potential influence of gut microbiota in ccRCC, but no obvious evidence indicates that a direct and exact relationship between gut microbiota and ccRCC tumorigenesis exists.

In this study, specific gut microbiota profiles in ccRCC patients compared to healthy controls were identified through16S rRNA gene sequencing. Additionally, metabolites and blood indices are detected and analyzed statistically. Bioinformatics analysis was also carried out to explore the potential mechanisms of ccRCC pathogenesis.

## RESULTS

### Variation of gut microbiota between the RCC group and the Control group

60 fecal samples were sequenced from the RCC group (n=30) and the Control group (n=30). Baseline clinical parameters of subjects were listed in Table 1. The RCC group and the Control group were similar in age, BMI and gender ratio (P>0.05).

**Table 1.** Baseline characteristics of all subjects. BMI, body mass index; WHO/ISUP, WHO/International Society of Urological Pathology; N/A, not applicable. Results were expressed as ratio or mean±s.d. Non-significant P>0.05.

| Characteristic | RCC Group (n=30) | Control Group (n=30) | P value |
| --- | --- | --- | --- |
| **Age (Year)**<br>(mean ± s.d) | 62±10 | 59±10 | 0.37 |
| **Gender**<br>(Ratio) | 73% (male) | 67% (male) | 0.14 |
| **BMI**<br>(mean ± s.d) | 25.85±3.42 | 24.23±2.97 | 0.06 |
| **Ethnicity – Han (China)**<br>(Ratio) | 30 (100%) | 30 (100%) | N/A |
| **Smoking**<br>(Ratio) | 37% (smoking) | 27% (smoking) | 0.42 |
| **Alcohol**<br>(Ratio) | 23% (alcohol) | 33% (alcohol) | 0.22 |
| **WHO /ISUP**<br>(Ratio) | 1: 9 (30%)<br>2: 14 (47%)<br>3: 6 (20%)<br>4: 1 (3%) | N/A | N/A |
| **TNM** | T1aN0M0 : 19 (63%)<br>T1bN0M0 : 7 (23%)<br>T2aN0M0 : 1 (3%)<br>T3aN0M0 : 3 (10%) | N/A | N/A |

BMI, body mass index; WHO/ISUP, WHO/International Society of Urological Pathology; N/A, not applicable. Results were expressed as ratio or mean ± s.d. Non-significant P>0.05.

A total of 1,800,840 valid sequences were obtained from 60 qualified fecal samples through samples sequencing and data processing. The Shannon and Simpson indices were shown in Figure 1A. Results indicated that there was no difference of the gut microbiota richness and evenness estimated by within-sample diversity in the RCC group compared to the Control group. More alpha diversity indices such as Chao 1, Observed species, Faith’s PD, Pielou’s evenness and Good’s coverage confirmed it (Figure S1). Beta-diversity analysis revealed there existed a significant difference in the overall structure of the gut microbiota between the RCC group and Control group (Figure 1B).

**Figure 1.**
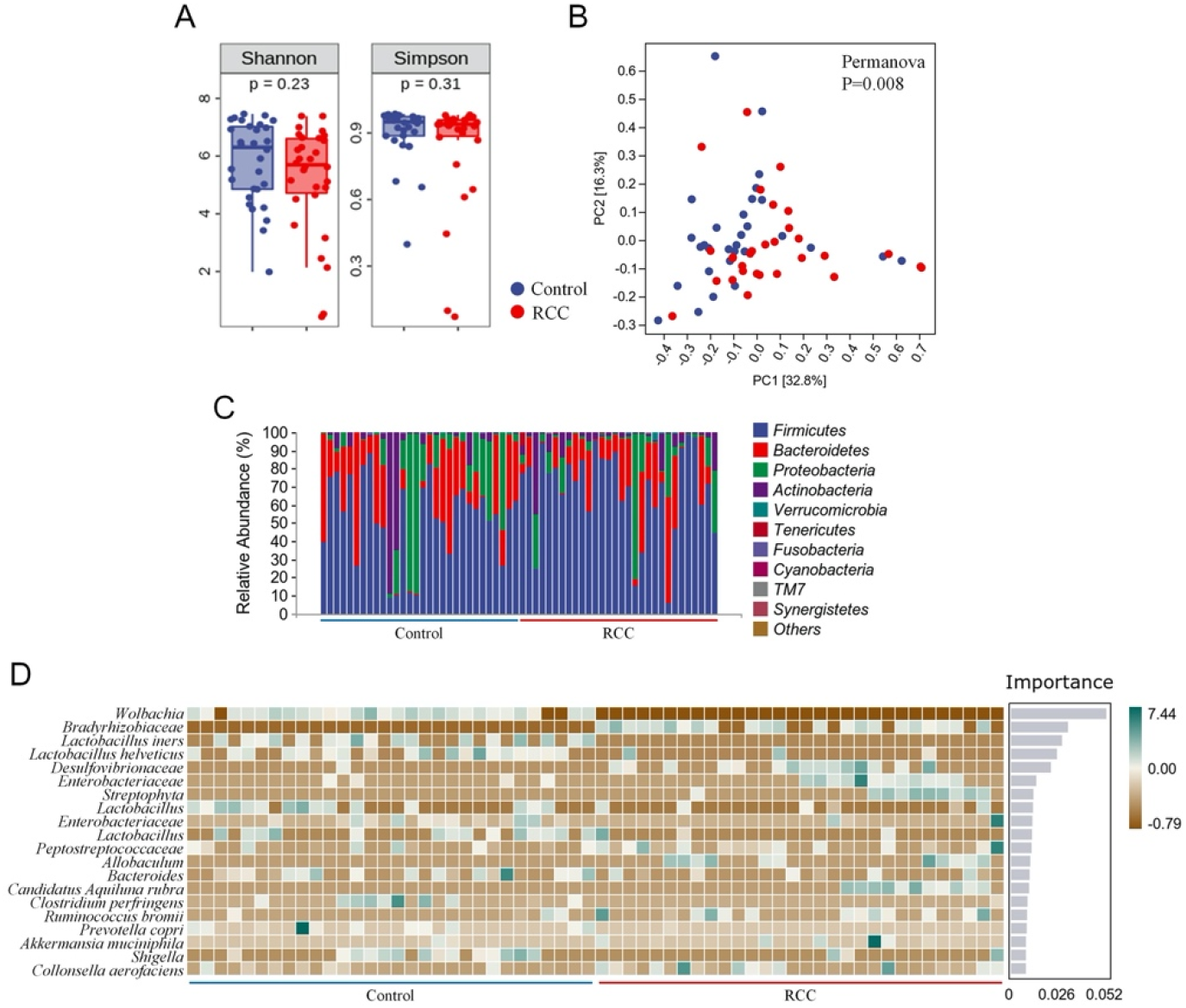
Characterization of gut microbiota were identified in the RCC and Control groups. (A) Alpha diversity of gut microbiota. The within-sample diversity of the RCC group and Control group were measured in terms of Shannon index and Simpson index. (B) Assessment of overall gut microbiota structures between the two groups using PERMANOVA analysis. PC1, principal coordinate 1; PC2, principal coordinate 2. (C) Relative abundance of gut microbiota in the two groups at phyla level. (D) Altered 20 genera significantly in the RCC group compared to Control group arranged based on the potential importance by Random Forest analysis. Green indicates positive correlation; Brown, negative correlation. Non-significant P>0.05.

To further clarify the features of gut microbiota in patients with ccRCC, we analyzed the relative abundance of specific microbiota, and classified them at different taxonomic levels. The abundance distribution of 10 main gut microbiota, including *Firmicutes, Bacteroidetes, Proteobacteria, Actinobacteria, Verrucomicrobia, Tenericutes, Fusobacteria, Cyanobacteria, TM7* and *Synergistetes* were described at phyla level (Figure 1C). Detailed taxonomic abundance difference of gut microbiota at family and phyla levels was shown according to the sequencing dataset (Figure S2). Moreover, there existed significant difference of 18 genera between the RCC group and the Control group by Random Forest analysis (Figure 1D). Results indicated that 11 species were enriched in ccRCC patients, including *Bradyrhizobiaceae, Desulfovibrionaceae,* two kinds of *Enterobacteriaceae, Streptophyta, Peptostreptococcaceae, Allobaculum, Candidatus Aquiluna rubra, Ruminococcus bromii, Akkermansia muciniphila* and *Collinsella aerofaciens,* while 9 species including *Wolbachia,* four kinds of *Lactobacillus, Bacteroides, Clostridium perfringens, Prevotella copri* and *Shigella,* were down-regulated notably. Moreover, it can be concluded from the results that *Wolbachia, Bradyrhizobiaceae* and *Lactobacillus iners* differed most typically based on the Importance. LEfSe was then applied on the analysis of significant microbial variations between the two groups, too (Figure S3). Our data preliminarily suggested that patients with ccRCC had alterations of specific bacterial functional species compared with those of the Control group.

### Identification of potential metabolic biomarkers for clear cell renal cell carcinoma

To pursue our primary hypothesis that changes of gut microbiota are associated with ccRCC tumorigenesis, we specifically examined the potential related differential genes involved in serum metabolites of the two groups. Except for the undefined superclass, the “organic acids and derivatives” accounted for 19.832% of all relevant metabolites with the largest proportion, according to the serum metabolomic analysis, followed by “lipids and lipid-like molecules” (Figure S4A). Volcano plots indicated that there were 256 altered metabolites in the RCC group compared to the Control group in positive ion mode and 102 in negative ion mode, and those metabolites were mostly classified as “organic acids and derivatives” and “lipids and lipid-like molecules” (Figure S4B and C).

A superior *OPLS-DA* model (Q^2^ > 0.5 both in positive and negative ion mode) was generated to analyze the difference of serum metabolomic profiles (Figure 2A and B). As shown in Figure 2C and D, 18 major metabolites altered significantly in positive ion mode were mostly classified as “organic acids and derivatives” based on superclass, and 7 major metabolites in negative ion mode were mostly classified as “lipids and lipid-like molecules”. Heatmap was also used to intuitively illustrate the abundance distribution of the 25 metabolites in all fecal samples screened by *OPLS-DA*, respectively, in positive ion mode (Figure 2E) and negative ion mode (Figure 2F). In general, 15 metabolites, such as taurine, were downregulated and 10 metabolites, for example, arachidonic acid, were upregulated in the RCC group.

**Figure 2.**
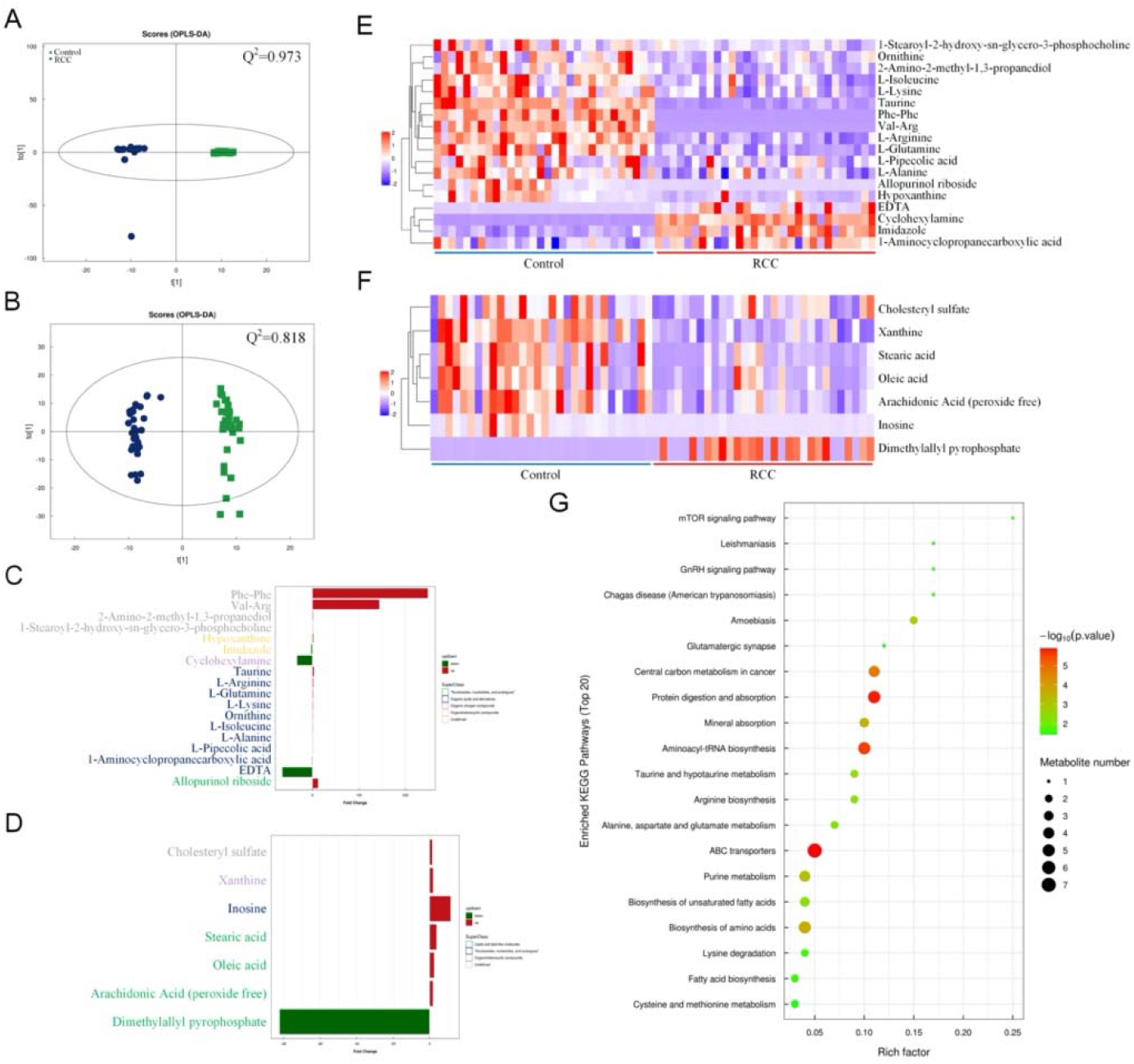
Serum metabolomics profiles of the RCC and Control groups were explored and visualized. Score plot of *OPLS-DA* in positive (A) and negative (B) ion mode, Q^2^ indicates the prediction ability of the model. Altered serum metabolites of ccRCC patients according to metabolomics analysis based on superclass in positive (C) and negative (D) ion mode. Abundance distribution of serum metabolites between the two groups analyzed by *OPLS-DA* and visualized by heatmaps in positive (E) and negative (F) ion mode, respectively. (G) Metabolic pathway impact map related to identified serum metabolites extracted from KEGG databases.

In order to explore the potential pathways by which the above altered metabolites function, a gene-KEGG pathways dot plot was applied (Figure 2G). The results showed that the identified metabolites involved 20 metabolic pathways, among which protein digestion and absorption, aminoacyl-tRNA biosynthesis and ATP binding cassette (ABC) transporter pathways were the most correlated ones. Meanwhile, a spearman analysis was conducted to investigate the patterns of interactions between metabolites that might play a role respectively in positive and negative ion mode (Figure S5). In the heatmap, the abundance of particular metabolites, such as taurine and arachidonic acid was further analyzed and demonstrated (Figure S6).

### Association between altered gut microbiota and clinical indices

Furthermore, clinical indices including blood routine, C-reactive protein, liver and kidney function and electrolyte analysis were also performed to evaluate the clinical status difference of the RCC group and the Control group (Figure 3 and S7). As shown in Figure 3, inflammation related indices such as WBC, CRP, neutrophil and MO, were significantly upregulated in the RCC group. We investigated whether the 20 microbial species differing between the RCC group and the Control group were correlated with well-established clinical indices and altered serum metabolites (Figure 4). Remarkably in the two groups, *Lactobacillus*, known as protective bacteria, was negatively correlated with inflammatory indices such as WBC, MO and CRP. Until now, we could draw the conclusion that inflammation induced by certain bacteria plays an essential role in ccRCC tumorigenesis.

**Figure 3.**
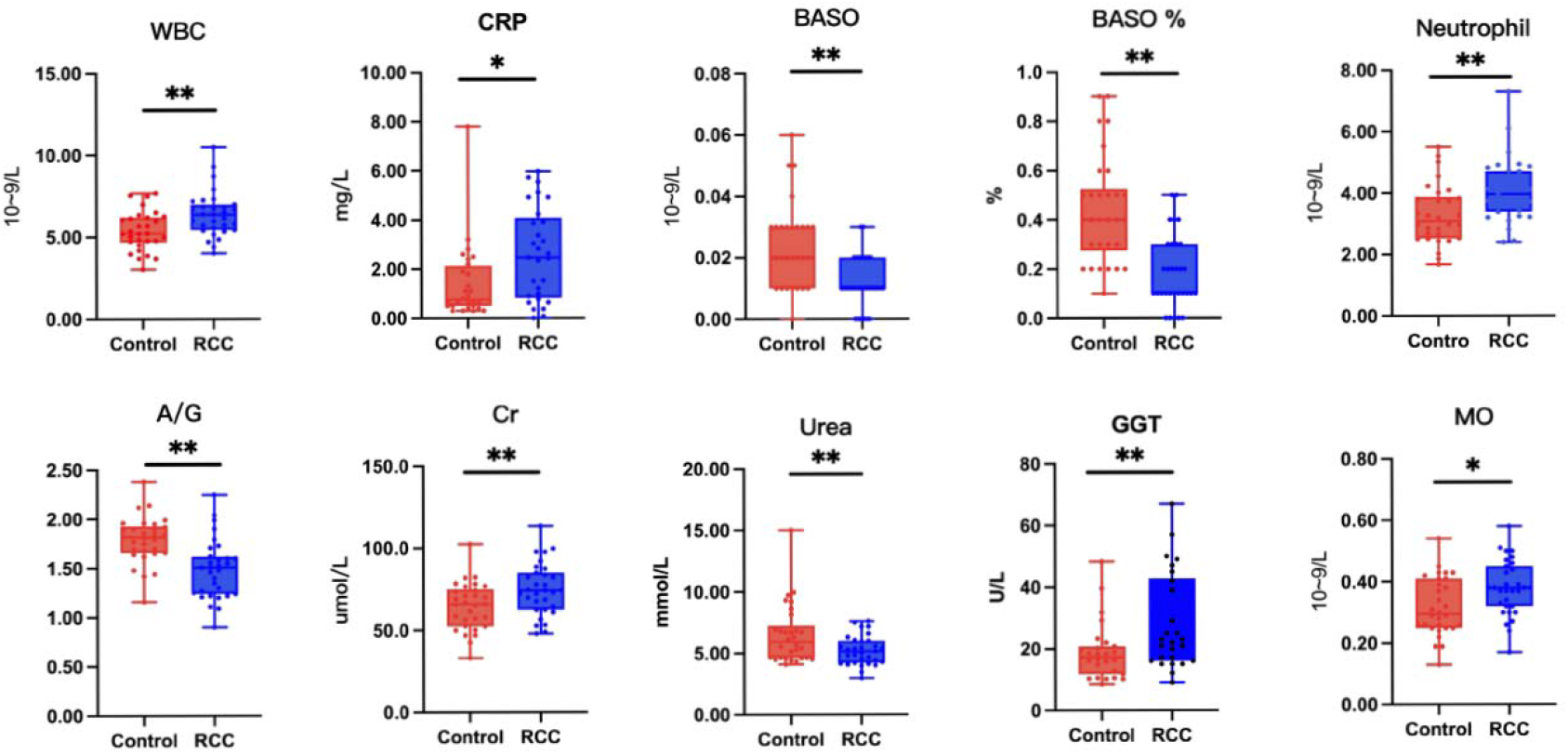
The differential levels of clinical indices were selectively detected between the RCC group and the Control group. WBC, white blood cell; CRP, C-reactive protein; BASO, basophil; A/G, albumin/globulin; Cr, creatinine; GGT, γ-glutamyl transpeptidase; MO, monocytes. *P < 0.05, **P < 0.01, ***P < 0.001.

**Figure 4.**
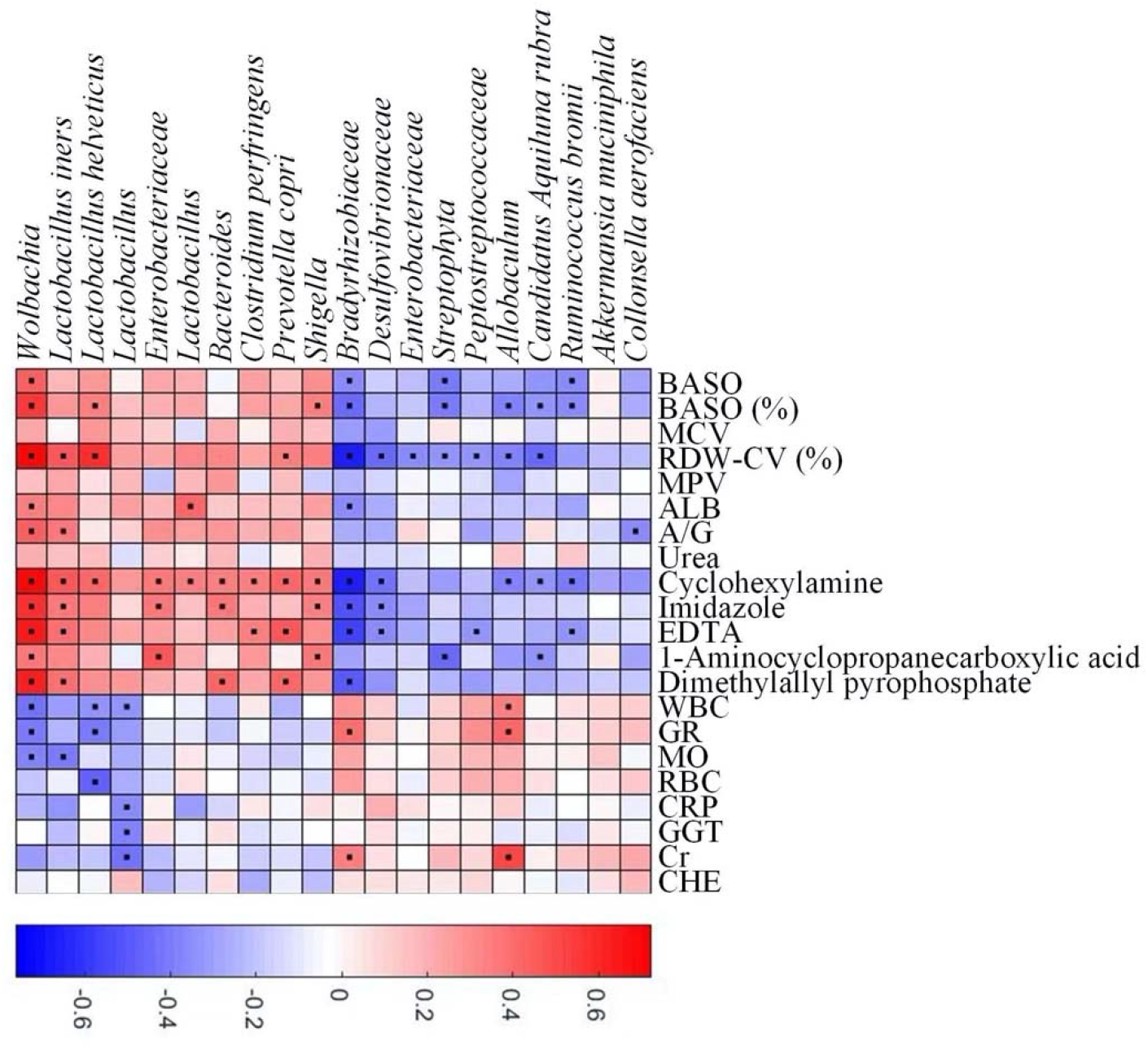
Correlation between altered genera and clinical indices of ccRCC patients. Red and blue color denoted positive and negative correlations, respectively. The black dots in the cells of the heatmap indicated the correlations were significant.

### Correlation and prognostic analysis of inflammation encoding genes and ccRCC development

Considering that the significant variation in a subset of gut microbiota might be closely associated with inflammation in the RCC group, we search for the possible link between inflammation encoding genes and ccRCC development. Result showed that 22 genes, including *Gabbr1* and etc., were identified as putative targets, which were overlapped across the univariate Cox regression and different expression analysis (Figure 5A, Table S1). We then estimated the influence of the 22 inflammatory factors on the prognosis of ccRCC (Figure 5B). Strikingly, *Gabbr1* is also a risk factor associated with poor prognosis of ccRCC.

**Figure 5.**
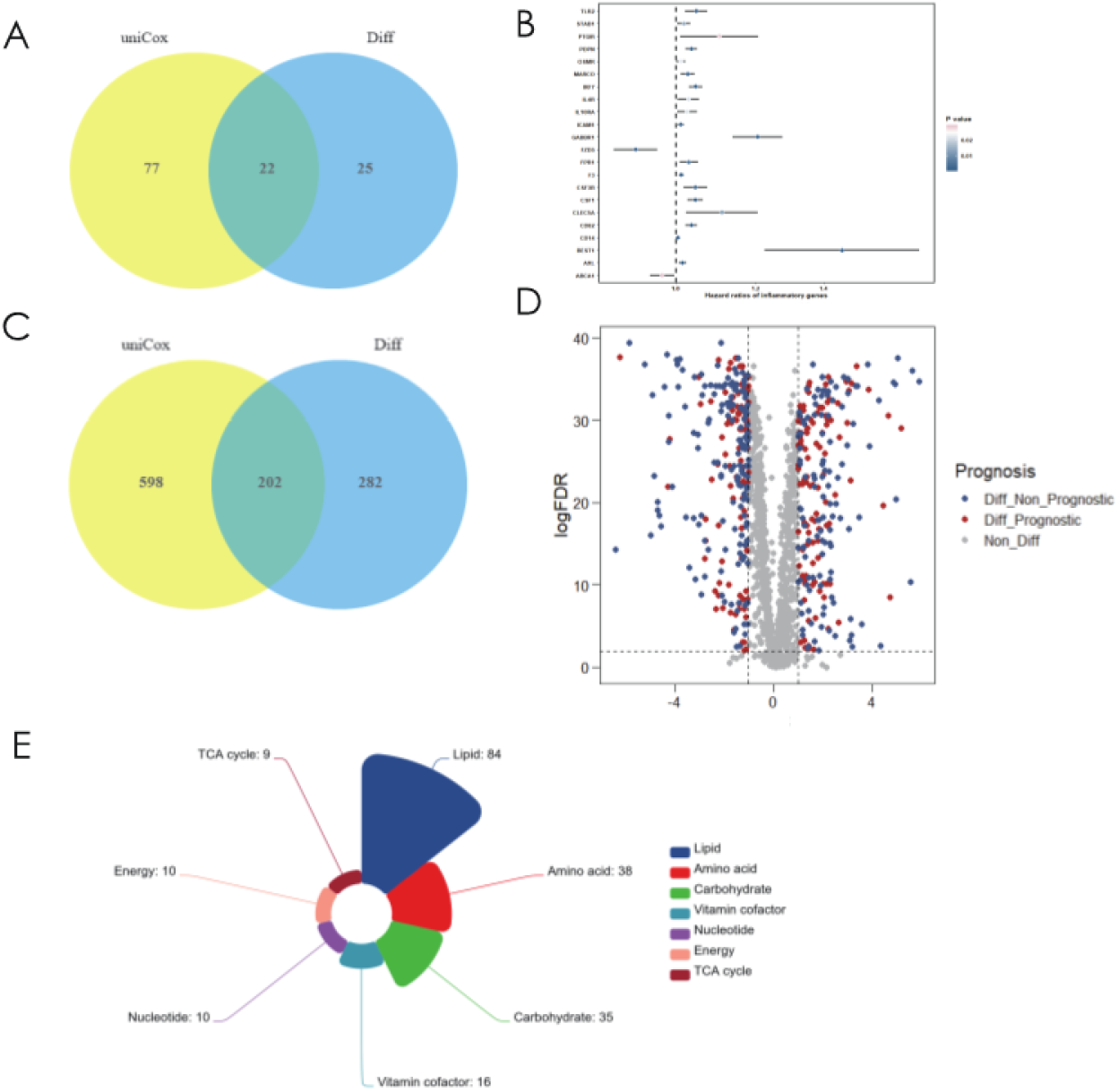
Bioinformatics analysis was applied to explore the association of gut microbiota and ccRCC tumorigenesis. (A) A total of 22 inflammatory transcripts were selected based on overlap between univariate Cox regression and different expression analysis; uniCox, univariate Cox regression analysis; Diff, different expression analysis. (B) The prognostic analyzes of 22 inflammatory transcripts by univariate Cox regression model. (C) A total of 202 metabolism-related transcripts were selected based on overlap between univariate Cox regression and different expression analysis. (D) Volcano plots selected 202 metabolism-related genes with significant prognosis from 484 different expression genes. (E) The 202 selected metabolism-related genes were clustered into 7 metabolism processes.

We also explored serum metabolites associated with ccRCC through *TCGA* database, in order to conduct further combined analysis with previous metabolomics results. The Venn diagram showed the overlap of 202 metabolism-related transcripts between the univariate Cox regression and different expression analysis (Figure 5C, Table S2). 202 metabolism-related genes were filtrated from different expression analysis with significant prognosis (Figure 5D). Finally, the 202 selected metabolism-related genes with significant different expression and prognosis were clustered into 7 metabolism processes, including lipid (42%), amino acid (19%), carbohydrate (17%), vitamin cofactor (8%), nucleotide (5%), energy (5%) and TCA cycle (4%) (Figure 5E). Results confirmed the crucial role of lipid metabolism in ccRCC tumorigenesis and development, which was almost consistent with our metabolomics results.

## DISCUSSION

The clinical features, early diagnosis and pathogenesis of renal cancer is still not fully understood (7, 17). Clear cell renal cell carcinoma, as the most common pathological type of renal cancer, deserves further attention and focus research (18). Recently, increasing evidence reveals that gut microbiota is closely associated with development and progression of multiply malignancies (19). In this study, we performed the 16S rRNA gene sequencing to comprehensively elucidate the variations of gut microbiota between the RCC group and the Control group. 11 species such as *Bradyrhizobiaceae* and *Desulfovibrionaceae* were enriched, and 9 species such as *Wolbachia* and four kinds of *Lactobacillus* were less abundant in the RCC group. Metabolomics analysis was applied to identify the difference of serum metabolites including taurine and arachidonic acid in the two groups. Gut microbiota of all subjects were analyzed based on taxonomic characterization and correlation with differential clinical indices of ccRCC patients. Our results indicated that specific component structures and functional patterns of gut microbiota and metabolites probably result in ccRCC tumorigenesis and development.

Diversity analyzes implied that though similar community richness and evenness are found in both groups, the overall structures between the two groups are different, where several altered bacterial species detected in the RCC group compared to the Control Group. Despite the perplexing taxonomy and synergy between the microbe, the effector microbiota that participated in or dominated the development of ccRCC may only be a small subset of them. Based on this, we particularly focused on *Desulfovibrionaceae* and *Lactobacillus. Desulfovibrionaceae* is a major producer of hydrogen sulfide, which acts as a poisonous regulator for mucosal function and intestinal environment. According to previous studies, *Desulfovibrionaceae* was considered to be a type of bacteria that posed health risk due to its positive correlation with intestinal and systematic inflammation (20, 21). Moreover, clinical data suggested that the presence of *Desulfovibrionaceae* was associated with chronic periodontitis, cell death, and inflammatory bowel diseases (22–24). With regard to *Lactobacillus,* it has been generally proved to play an essential role in maintaining homeostasis. *Lactobacillus* functions positively in immune regulation, which can significantly promote antibody production, activate and enhance macrophages, so as to improve the disease resistance of the body (25, 26). The occurrence of multiple cancers including ccRCC has been proved to be closely related to inflammatory factors and the pathways they dominate (27, 28). Other altered microbiota such as *Enterobacteriaceae, Streptophyta* and *Peptostreptococcaceae* have also been proved to be linked to some urinary system diseases associated with inflammation such as prostatitis and prostate cancer (29, 30). In summary, these results were consistent with the hypothesis that ccRCC tumorigenesis was accompanied by variation in specific gut microbiota profiles.

In order to verify whether the alteration of gut microbiota could cause changes in host metabolism, we further conducted a metabolomic analysis to identify the differential serum metabolites. 358 altered metabolites in total were detected in the RCC group compared to the Control group, both in positive and negative ion modes. By cluster analysis, we found that the altered metabolites are mainly classified as organic acids and derivatives or lipids and lipid-like molecules. These results indicated that the abnormality of amino acid metabolism and lipid metabolism might be involved in ccRCC tumorigenesis.

Notably, we specifically focused on taurine from 25 altered metabolites screened by *OPLS-DA*. Kidney is the main excretion and content regulating organ of taurine, and taurine is consumed by *Desulfovibrionaceae* and released by *Lactobacillus* in gastrointestinal tract. As a source of sulfur-containing substances, taurine is required by *Desulfovibrionaceae* as a growth stimulator that is reduced to H2S to some extent, which in turn induces intestinal pathology (31). *Lactobacillus*, one of the crucial components of probiotics, can promote the activity of bile salt hydrolase to increase the abundance of taurine in the intestine, thereby stimulating tight junctions and reducing inflammatory responses (32).

Taurine has always been considered as a non-functional metabolite of sulfur-containing amino acids, widely distributed in the human body in the form of free amino acids. However, taurine is involved in cell protection, renal insufficiency, and abnormal renal development. It has a wide range of biological functions such as attenuating inflammation and oxidative stress-mediated injuries, lipid metabolism regulation and immune enhancement (33–35). Excessive reactive oxygen species (ROS) formation is one of the determinants in oncogenesis and is no exception in the development of ccRCC (36). Taurine can prevent the accumulation of ROS in tumor cells, enhance immune surveillance, as well as confer its preventive effect on cancer cells. In addition, taurine induces the apoptosis of tumor cells by up-regulating the expression of the p53 transcription factor, down-regulating the expression of B-cell lymphoma 2 (BCL-2) or inactivating the protein kinase B (Akt) signaling pathway and etc., thereby may inhibit ccRCC progression (37). Based on the above findings, we could assume that the decrease of serum taurine was relevant for the increased *Desulfovibrionaceae* and reduced *Lactobacillus* in gut microbe community, which in turn contributed to ccRCC tumorigenesis probably.

Chronic inflammation in the tumor microenvironment has been shown to contribute to ccRCC progression previously. In ccRCC tumors, IL-1β, IL-6 and TNF are essential inflammatory mediators and high serum levels of these cytokines are associated with tumor progress, by inducing the activation of endothelial cells or promoting angiogenesis (38). Recently, alteration of gut microbiota is regarded as a trigger for systemic inflammation especially in metabolic diseases, as well as tumorigenesis by modulate the secretion of inflammation-related products such as SCFA and TMAO (39, 40).

To shed light on the genetic signature changes that gut microbiota possibly functioned during ccRCC tumorigenesis, we performed a bioinformatics analysis of ccRCC-related metabolites and inflammation in the *TCGA* database. Combined with cluster analysis, we established a robust association between altered microbiota and ccRCC. In our study, correlation analysis indicated that inflammation related clinical indices were significantly upregulated in the RCC group even though the values were in the normal range. Consequently, we screened out 22 inflammatory genes such as *Gabbr1,* by univariate Cox regression and different expression analysis in the *TCGA* database. *Gabbr1* was classical macrophage-related genes which have been confirmed to be the inextricable link between inflammation and ccRCC (41, 42). Other overlapped genes, such as FZD5 encoding WNT signal molecular and STAB1, which are known as a scavenger receptor also participate in carcinogenesis, and their potential functions in ccRCC need to be further revealed (43, 44). We also found that the altered metabolites mostly participated in protein digestion and absorption, aminoacyl-tRNA biosynthesis and ABC transporter pathways. In addition, we performed prognostic analysis on 202 selected metabolites associated with ccRCC to refine the above conjecture.

These findings provide evidence for the hypothesis that the alterations of gut microbial composition are associated with ccRCC. Reduction of protective bacteria, proliferation of sulfide-degrading bacteria Desulfovibrionaceae, reduction of taurine, and enrichment of macrophage related genes more prone to ccRCC tumorigenesis. Systemic low-grade inflammation as well as abnormal lipid metabolism and amino acid metabolism may be the functional bridge between dysbiosis and ccRCC.

Several limitations should be taken into account in our study. First, how changes in gut microbiota regulate the pathogenesis of ccRCC or whether changes in gut microbiota are caused by ccRCC remain to be further explored. Second, fecal samples were not subjected to additional metabolomic analysis, so the synchronization of metabolite changes in feces and serum could not be determined.

In future studies, we will validate the effects of taurine and the two species on ccRCC using fecal microbiota transplantation (FMT) in vivo by animal experiments. Although clinical indicators suggested that the altered gut microbiota were related to inflammation, the detection of specific inflammatory factors, such as IL-4 and IL-10, and the exploration of more in-depth molecular mechanisms still needs to be completed.

## CONCLUSION

In conclusion, our study suggests a new avenue that links the alteration of gut microbial composition and function, and the systemic inflammatory and metabolic state of ccRCC. The protective bacteria *Lactobacillus* and sulfide-degrading bacteria *Desulfovibrionaceae* are the main effective microbiota of ccRCC, and taurine and inflammatory factors may be the mediators between Pathogenic microbiota and ccRCC.

## MATERIALS AND METHODS

### Subjects

A clinical trial with 30 ccRCC patients (RCC group) and 30 healthy controls (Control group) was conducted. All ccRCC patients in the RCC group were diagnosed exactly by pathological examinations after surgery, and all healthy controls were recruited from Beijing Friendship Hospital, Capital Medical University (Beijing, China). The healthy controls at this stage were free of renal cell carcinoma by medical examinations and had no history before enrollment. All subjects were Chinese Han ethnic population to ensure their living and eating habits were basically similar. All experimental subjects conformed to the following standards. ***Inclusion Criteria*:** (1) no infectious disease or abnormal renal function, no neurological symptoms, no upper gastrointestinal disease, (2) performance status (PS) score ranging from 0 to 1.5, proper functioning of major organs, no other active malignancies (except non-melanoma skin cancer), (3) no antibiotics, probiotics, vitamins, minerals, NSAIDs, prokinetics, bismuth, antacids, H2-receptor antagonists, omeprazole, Sucralfate or misoprostol intake within the past 4 weeks, no recent hormone therapy taken, (4) no history of gastroduodenal ulcer or major gastrointestinal surgery. ***Exclusion criteria*:** (1) Patients with gastrointestinal discomfort such as acid reflux and nausea, diabetes or other serious systemic diseases (2) HIV-infected or hepatitis B surface antigen positive patients, (3) patients with incomplete heart function, unstable angina pectoris or recent (within 6 months) myocardial infarction, (4) patients with history of major organ transplant, radiotherapy or chemotherapy, (5) females who are pregnant or breast-feeding.

The study was approved by the Ethics Committee of Beijing Friendship Hospital Affiliated to Capital Medical University (Beijing, China). All subjects have signed the informed consent.

### Fecal samples collection and blood index detection

Patients in the RCC group were all initially diagnosed as renal carcinoma by CT/MRI scan. After being informed about the purpose and procedures of the study, sufficient, middle and internal fecal samples were freshly collected from both of the RCC Group and Control Group by the laboratory personnel with special fecal sampling tubes, and samples were transported to the laboratory on ice within one hour (Xing Kang Medical, Jiangsu). Fecal samples of patients in the RCC Group were all collected before surgery, and urine and urinal pollution to the fecal samples were avoided during collection processes. Samples were stored at −80 °C until further processing.

A fasting blood sample of all subjects were taken before receiving any medical treatments or surgery. Serum specimens were separated by centrifugation for a thorough analysis, including blood routine test, C-reactive protein test, liver and kidney function test, as well as electrolyte detection.

Note that the fecal and blood sample used for experiments and analysis are from patients in RCC group that have postoperative pathological diagnosis confirmation of ccRCC.

### Microbial DNA extraction and sequencing

Total microbial DNA was extracted from the fecal samples using the *OMEGA Soil DNA Kit* (Omega BioTek, Norcross, GA, USA) according to the manufacturer’s instructions. The DNA concentration were evaluated in a *NanoDrop ND-1000* spectrophotometer (Thermo Fisher Scientific, Waltham, MA, USA) respectively and all DNA samples were stored at −80 °C prior to further analysis. The A260:A280 ratios were ranging from 1.8 to 2.0.

PCR amplification of the microbial 16S rRNA V3–V4 hypervariable regions was performed. PCR primers of the V3-V4 hypervariable region were designed as previously described (45). The forward primer used was 5’-TCGTCGGCAGC GTCAGATGTGTATAAGAGACAGCCTACGGGNGGCWGCAG-3’, and the reverse primer was 5’-GTCTCGTGGGCTCGGAGATGTGTATAAGAGACAGGACTACHVGG GTATCTAATCC-3’. Amplification was performed on an ABI 9600 instrument according to the instruction successively with a denaturation at 94 °C for 4 min, 25 cycles of denaturation at 94 °C for 45 s, annealing at 60 °C for 45s, elongation at 72 °C for 45 s and a final extension cycle at 72 °C for 8 min.

Gut microbial 16S rRNA gene V3-V4 region was sequenced using *Illumina Miseq* platform as previously described (45). The high-quality data was obtained after quality control processing. Bioinformatics analysis of sequencing data was performed using *QIIME2* (46).

After removal of primers with *Cutadapt*, the sequences were denoised, merged and chimera were removed with *DADA2*. The remaining high-quality sequences were clustered into amplicon sequence variants (ASVs). A representative sequence from each ASV was assigned taxonomically against the *SILVA* Database. Next, an original ASV table was created which contained a readable matrix of the ASV abundance for each sample. Alpha diversity based on Shannon, Simpson, Chao 1, Observed species, Faith’s PD, Pielou’s evenness, and Good’s coverage were estimated. Beta-diversity was calculated based on Euclidean distance, and the significance of difference between the two groups was assessed by permutational multivariate analysis of variance (PERMANOVA) test in *MATLAB R2018b* (The MathWorks, Inc., Natick, MA, USA). Random Forest and linear discriminant analysis effect size (LEfSe) were contrasted based on the relative abundance of ASVs. The top 20 most important ASVs were shown in Random Forest. The ASVs with significant changes were found out in LEfSe when alpha value of the factorial Kruskal-Wallis test was <0.05 and the logarithmic LDA score was >2.0. The associations between the differential ASVs and the blood biochemical and metabolic indexes were performed by Spearman correlation.

### Blood metabolites analysis

Blood samples were collected in Vacutainer tubes containing the chelating agent ethylene diamine tetraacetic acid (EDTA). the samples were then centrifuged at 1500g, 4°C for 15min. Plasma samples were stored at −80 °C prior to further analysis.

200 ul of plasma samples were mixed with 400 ul of cold methanol/acetonitrile (1:1, v/v) to remove the protein. The mixture was centrifuged 14000 g, 4 °C for 15 min. and the supernatants were dried in a vacuum centrifuge. The samples were re-dissolved in 100 ul acetonitrile/water (1:1, v/v) solvent.

Analysis was performed using an *UHPLC* (1290 Infinity LC, Agilent Technologies) coupled to a quadrupole time-of-flight (AB Sciex TripleTOF 6600). Samples were analyzed by HILIC separation. A 2.1mm*100mm ACQUIY UPLC HSS T3 1.8um column (waters, Ireland) was used. In ESI positive mode, the mobile phase contained A=water with 0.1% formic acid and B=acetonitrile with 0.1% formic acid; and in ESI negative mode, the mobile phase contained A=0.5mM ammonium fluoride in water and B=acetonitrile. The gradient was 1% B for 1.5 min and was linearly increased to 99% in 11.5 min and kept for 3.5 min. then it was reduced to 1% in 0.1 min and a 3.4 min of re-equilibration period was employed. The gradients were at a flow rate of 0.3 mL/min, and the column temperatures were kept constant at 25°C. 2rewq 2ul aliquot of each sample was injected.

The ESI source conditions were set as follows: Ion Source Gas1 (Gas1): 60, Ion Source Gas2 (Gas2): 60, curtain gas (CUR): 30, source temperature: 600°C, IonSpray Voltage Floating (ISVF): ±5500V; TOF MS scan m/z range: 60-1000 Da, product ion scan accumulation time: 0.20 s/spectra; TOF MS/MS scan m/z range: 25-1000 Da, product ion scan accumulation time: 0.05 s/spectra. The product ion scan was acquired using information dependent acquisition (IDA) with high sensitivity mode selected. The parameters were set as follows: collision energy (CE): 35V±15eV; declustering potential (DP): ±60V; exclude isotopes within 4 Da; candidate ions to monitor per cycle: 10.

The raw MS data were converted from wiff.scan files to MzXML files using ProteoWizard MSConvert, then data were imported into freely available SCMS software for peak picking and grouping. Collection of Algorithms of Metabolite pRofile Annotation (CAMERA) were used for annotation of isotopes and adducts. Compound identification of metabolites was performed by comparing of accuracy m/z value (<25 ppm), and MS/MS spectra with an in-house database established with available authentic standard.

After normalized to total peak intensity, the data were then analyzed suing R package (v3.2.0) (47). Univariate analysis and orthogonal partial least-squares discriminant analysis (OPLS-DA) were performed to find out the differential metabolites between the two groups. The 7-fold cross-validation and response permutation testing was used to evaluate the robustness of the model. The variable importance in the projection (VIP) value of each variable in the OPLS-DA model was calculated to indicate its contribution to the classification. Metabolites with the VIP value >1 was further applied to Student’s t-test at univariate level to measure the significance of each metabolite, the p values less than 0.05 were considered as statistically significant. After combining the differential metabolites screened by positive and negative ion modes, the KEGG pathway was annotated and analyzed.

### Metabolic subtypes and inflammatory analysis

In brief, public gene expression data of ccRCC patients was downloaded from the Cancer Genome Atlas (*TCGA*) database. A well-established metabolic signature gene set was adopted in a previous study (48), which included genes for amino acid metabolism (348 genes), carbohydrate metabolism (286 genes), energy metabolism (110 genes), lipid metabolism (766 genes), nucleotide metabolism (90 genes), tricarboxylic acid cycle (TCA cycle, 148 genes), and vitamin cofactor metabolism (168 genes). A total number of 200 genes that related to inflammatory response were also collected. Differentially expression genes between tumor and normal tissues were detected by “limma” R package with fold change > 2 and false discovery rate < 0.05. In addition, we performed the prognostic analysis for selected genes by using univariate Cox regression model. Finally, all the data was visualized by different R packages.

### Data availability

The raw sequencing data are available at the Sequence Read Archive (SRA) of the National Center for Biotechnology Information (NCBI) under accession number PRJNA908738.

## SUPPLEMENTAL MATERIAL

Supplemental material is available online only.

FIG S1, TIF file, 1.3MB.

FIG S2, TIF file, 3.6MB.

FIG S3, TIF file, 1.3MB.

FIG S4, TIF file, 2.2MB.

FIG S5, TIF file, 1.8MB.

FIG S6, TIF file, 0.7MB.

FIG S7, TIF file, 1.6MB.

TABLE S1, PDF file, 0.1MB.

TABLE S2, PDF file, o.4MB.

## ACKNOWLEDGEMENTS

This work was supported by grants from the National Natural Scientific Foundation of China (Grant No. 82002711), Beijing Municipal Administration of Hospitals’ Youth Programme (Code: QML20200105), Training Fund for Open Projects at Clinical Institutes and Departments of Capital Medical University (CCMU2022ZKYXY019).

---

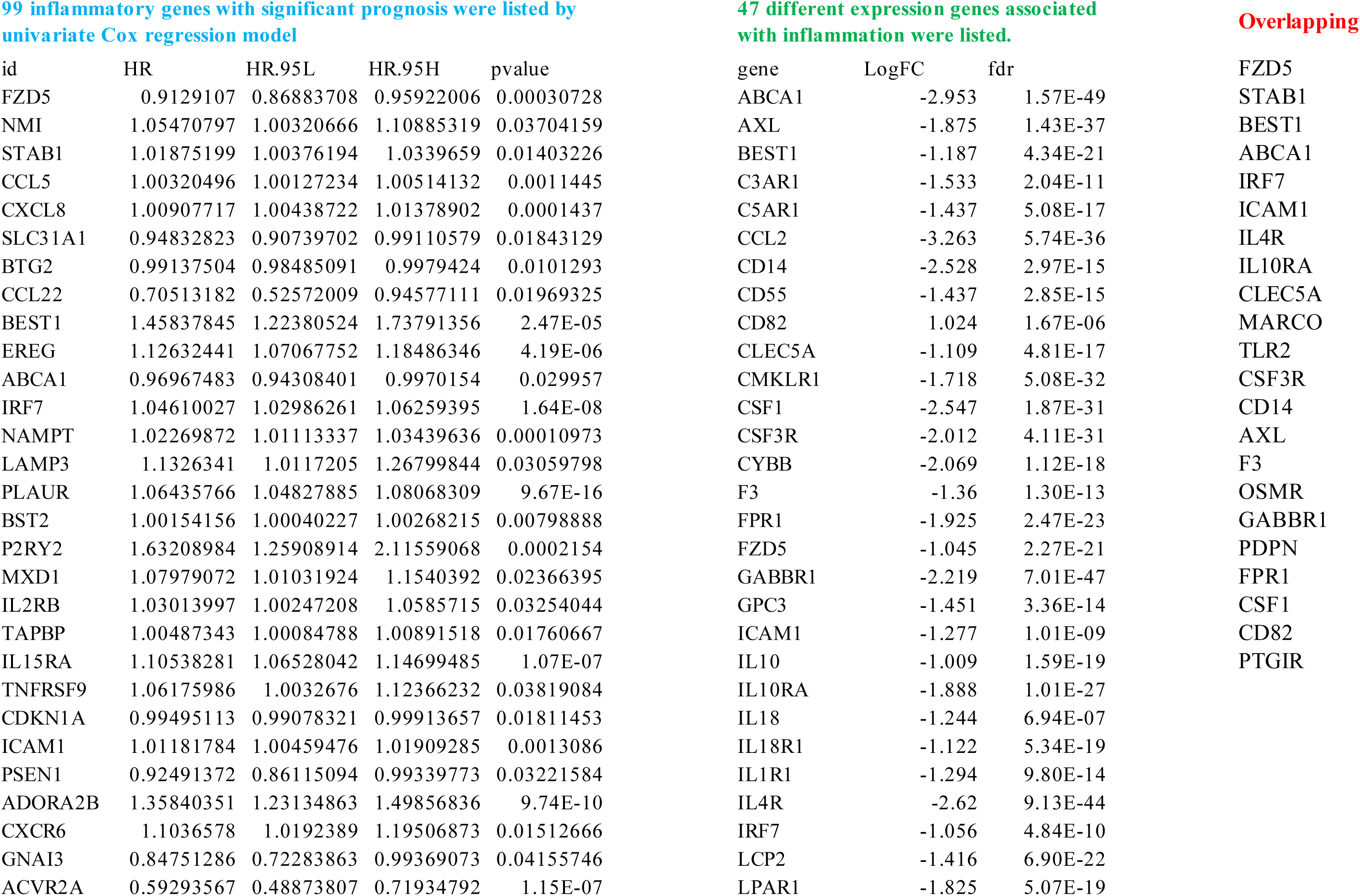
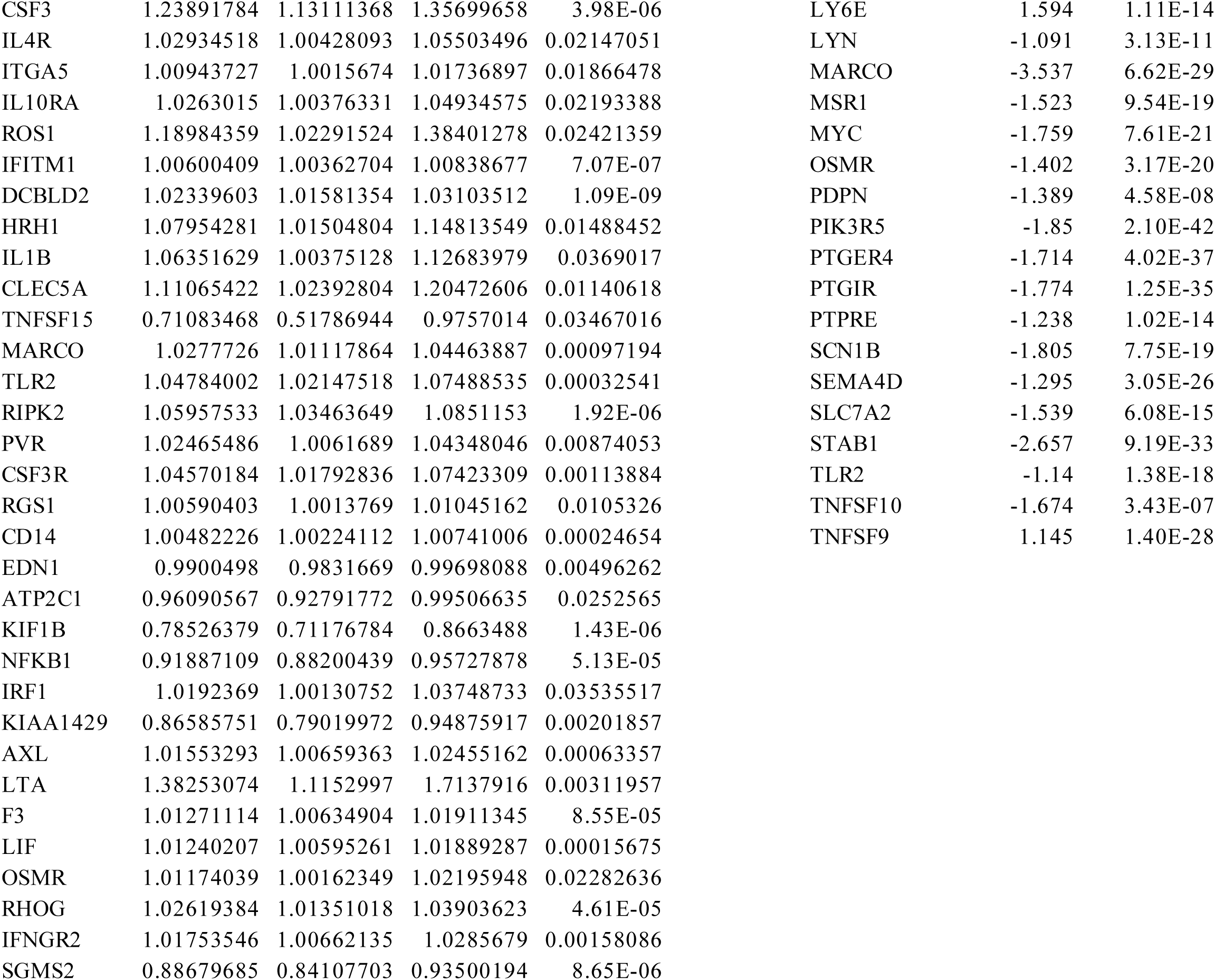
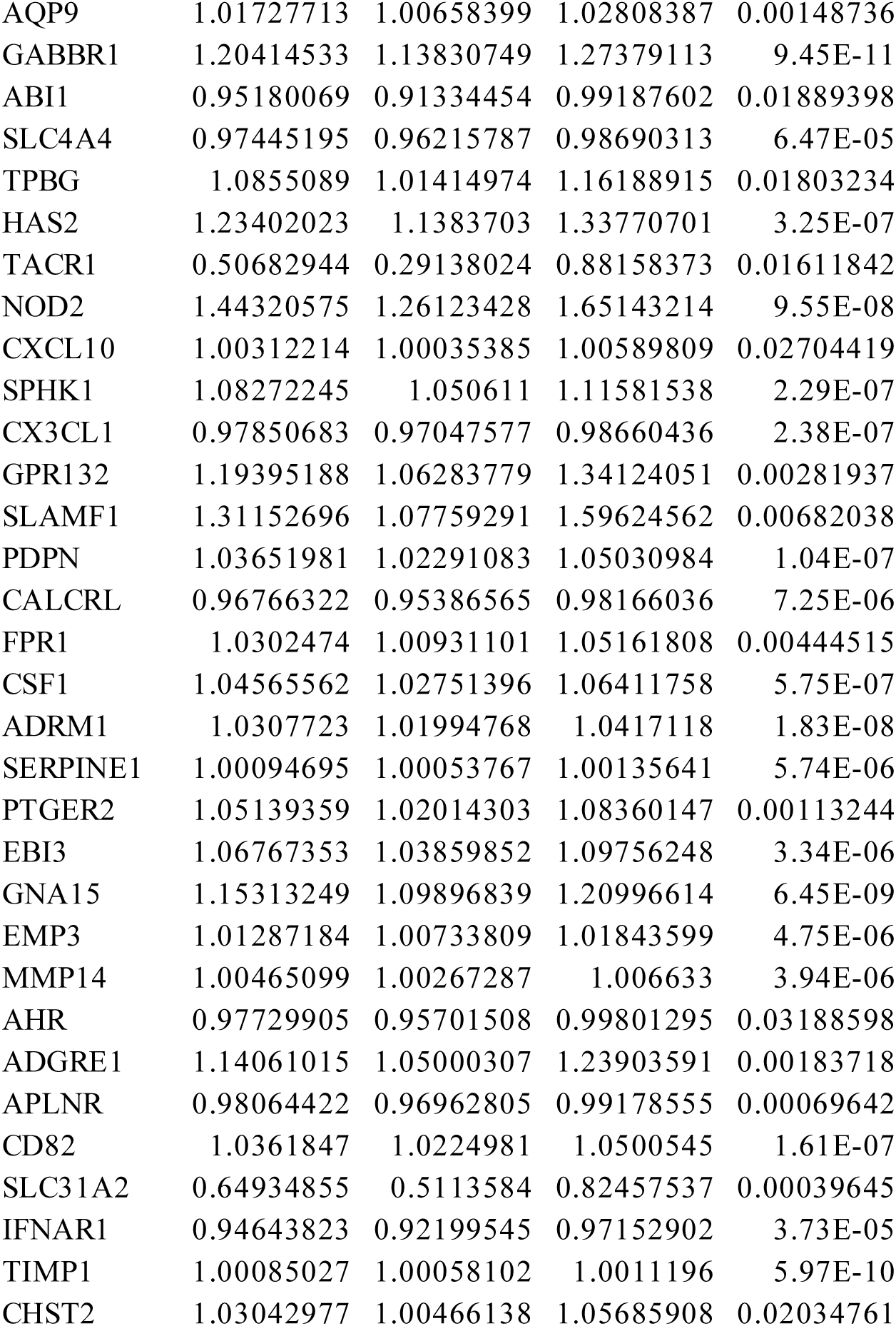
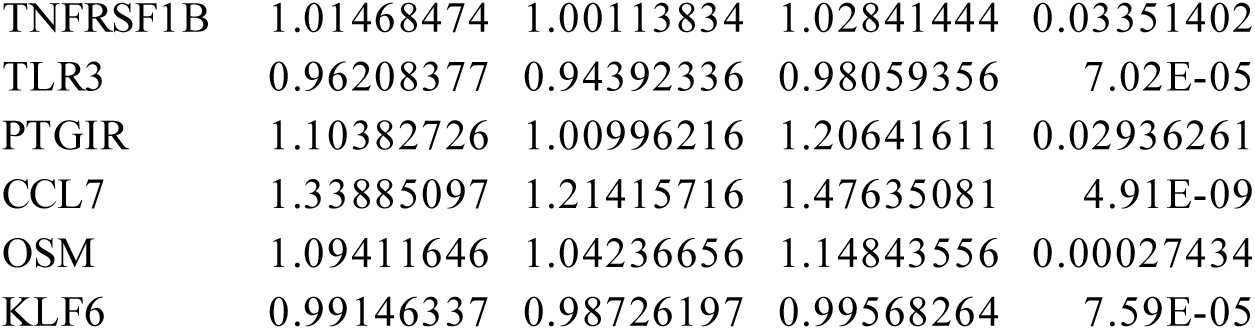

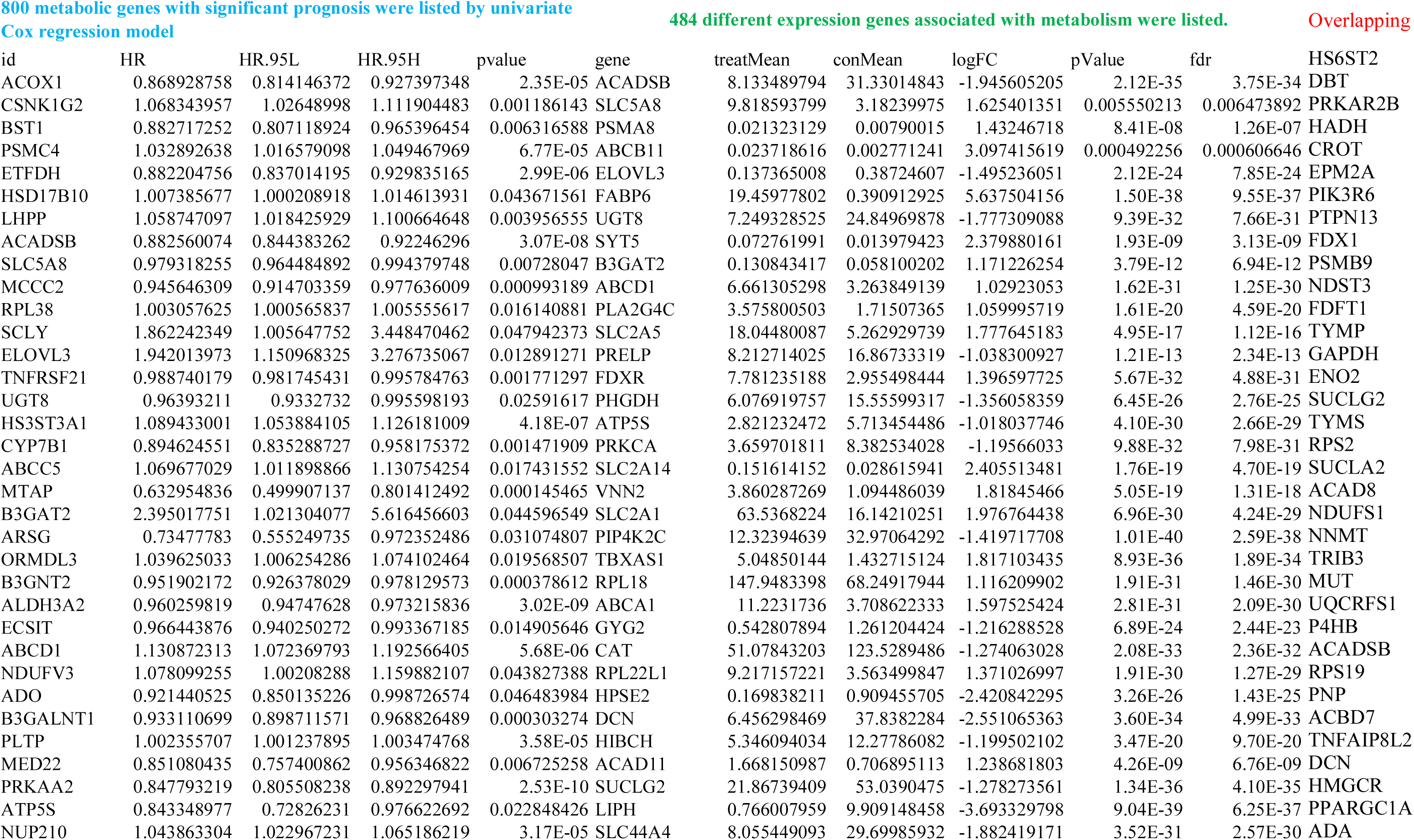
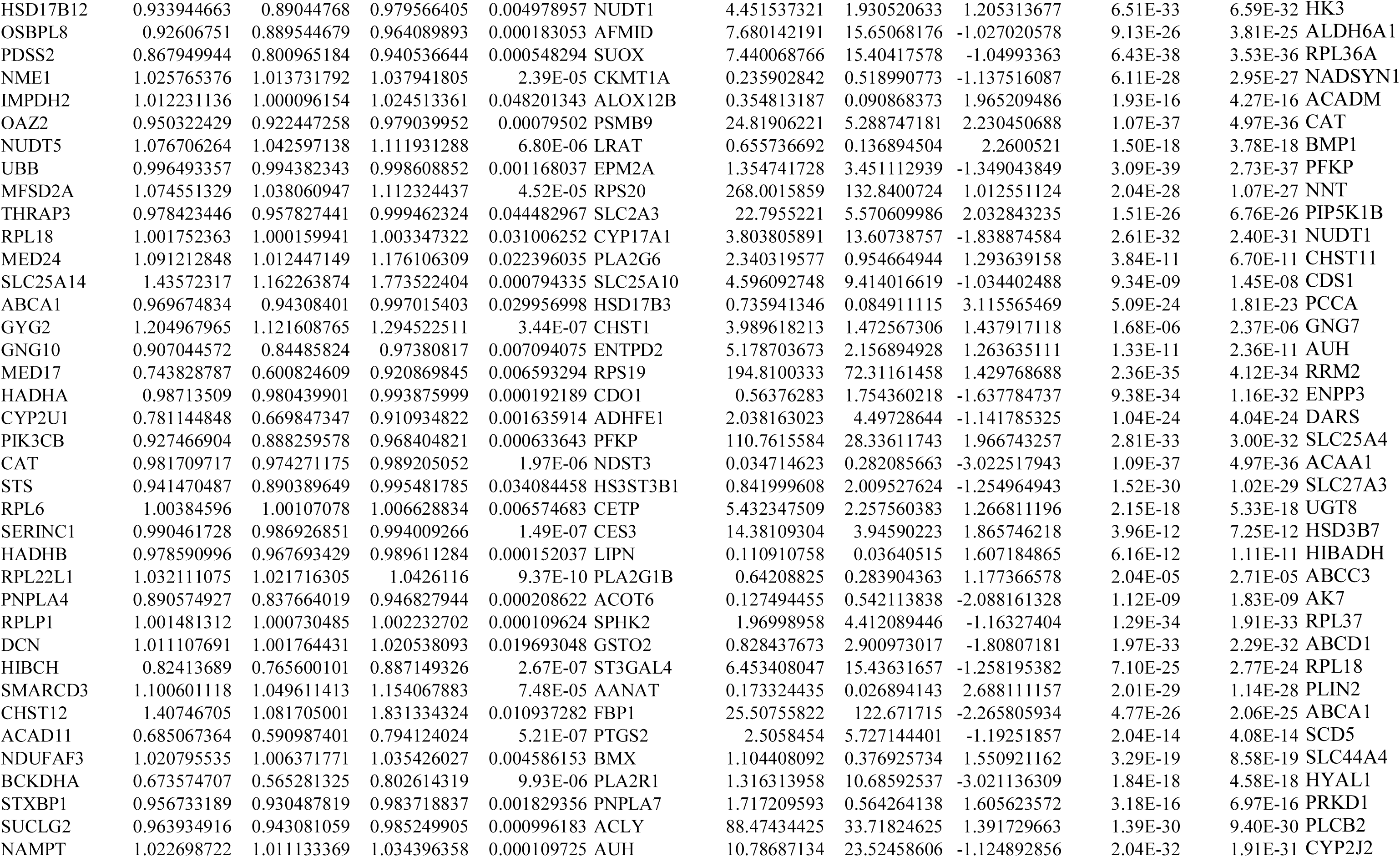
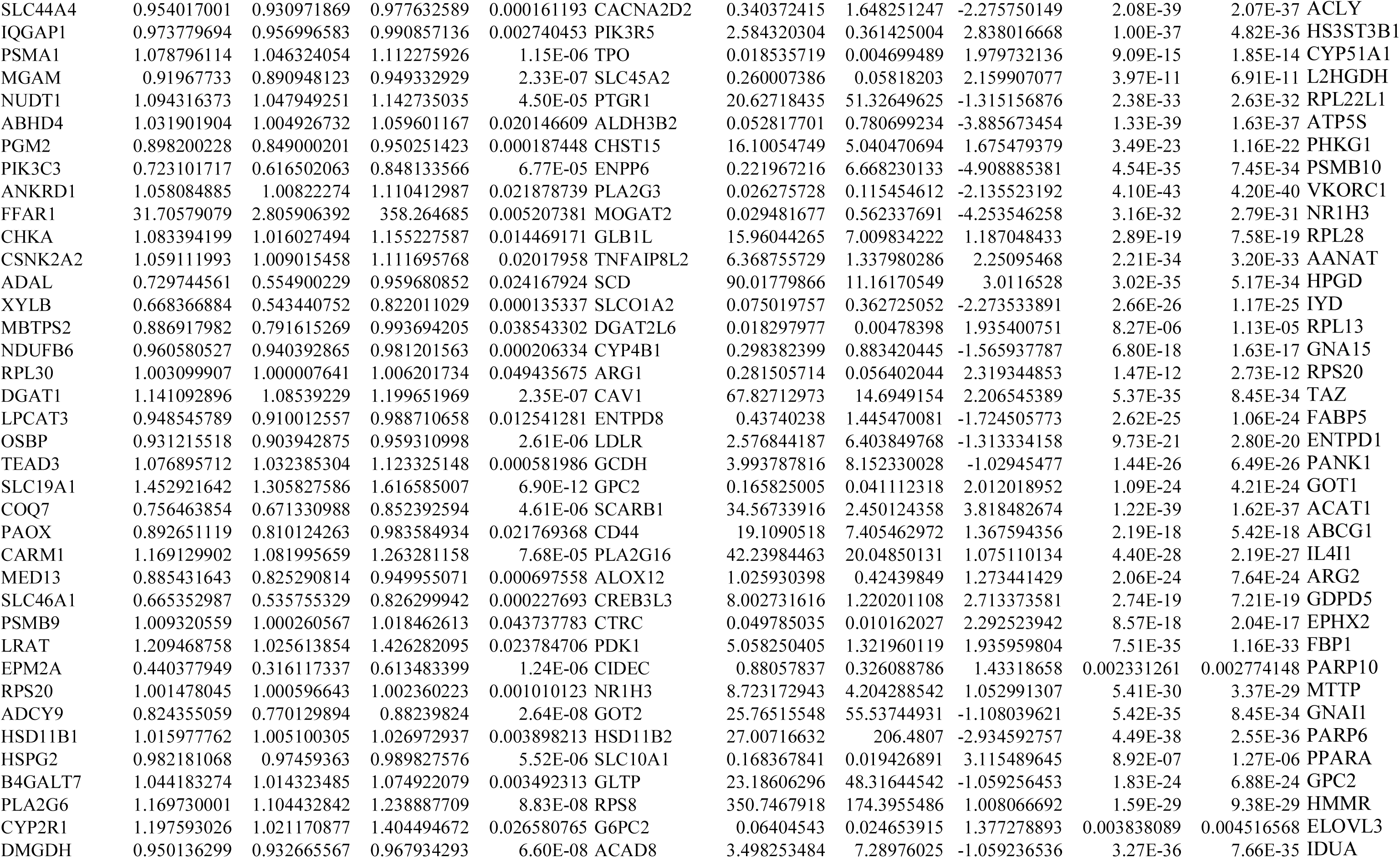
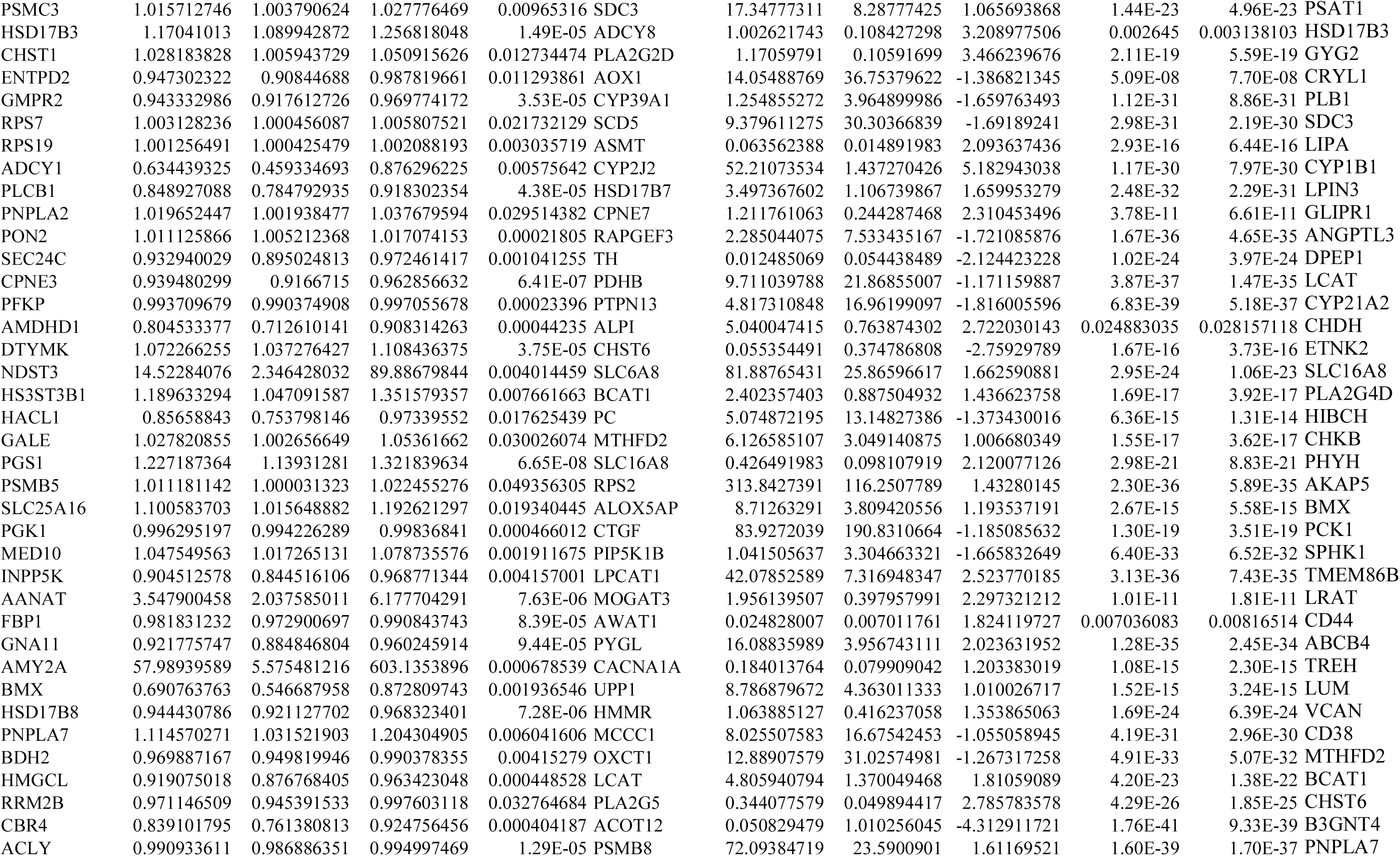
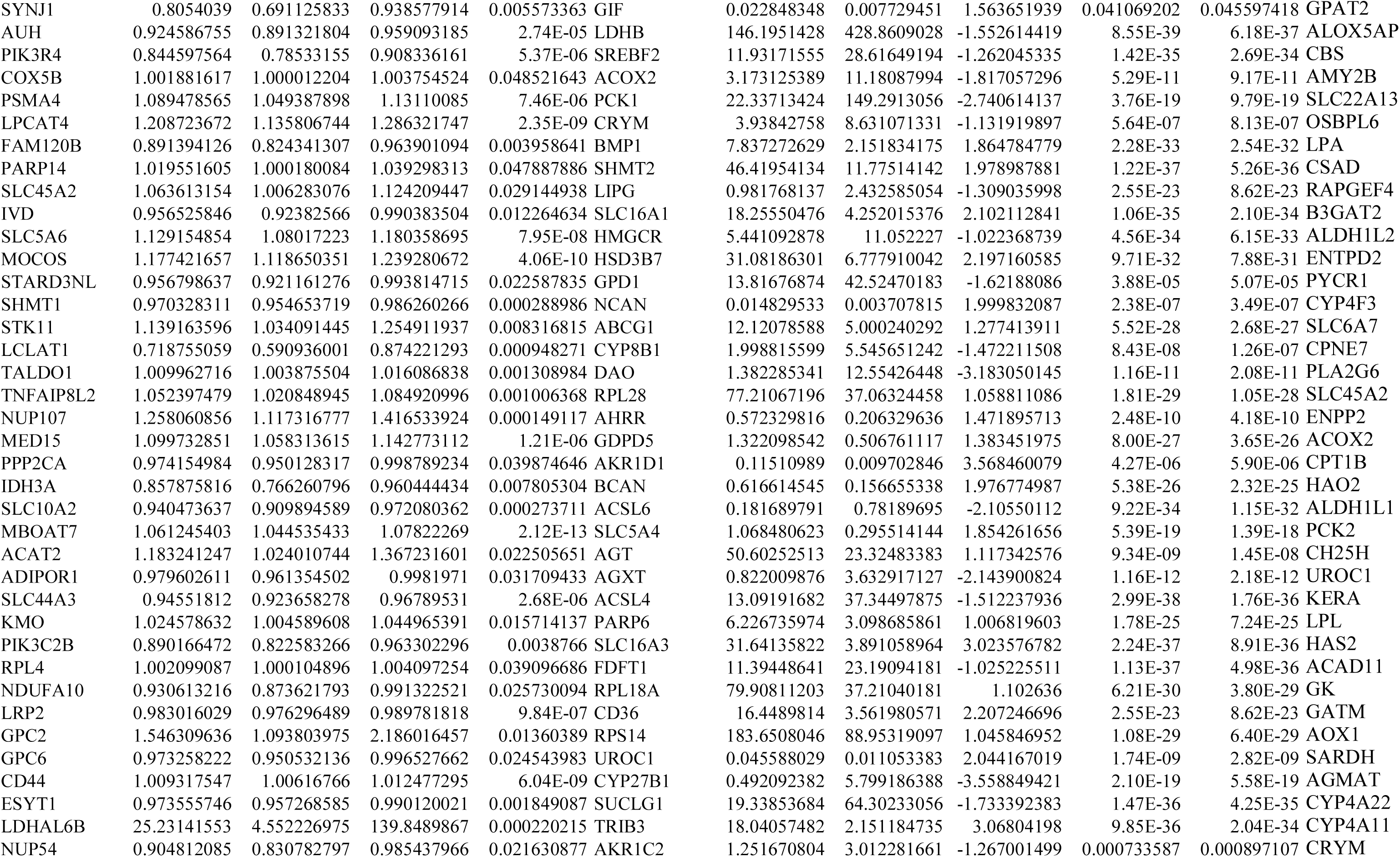
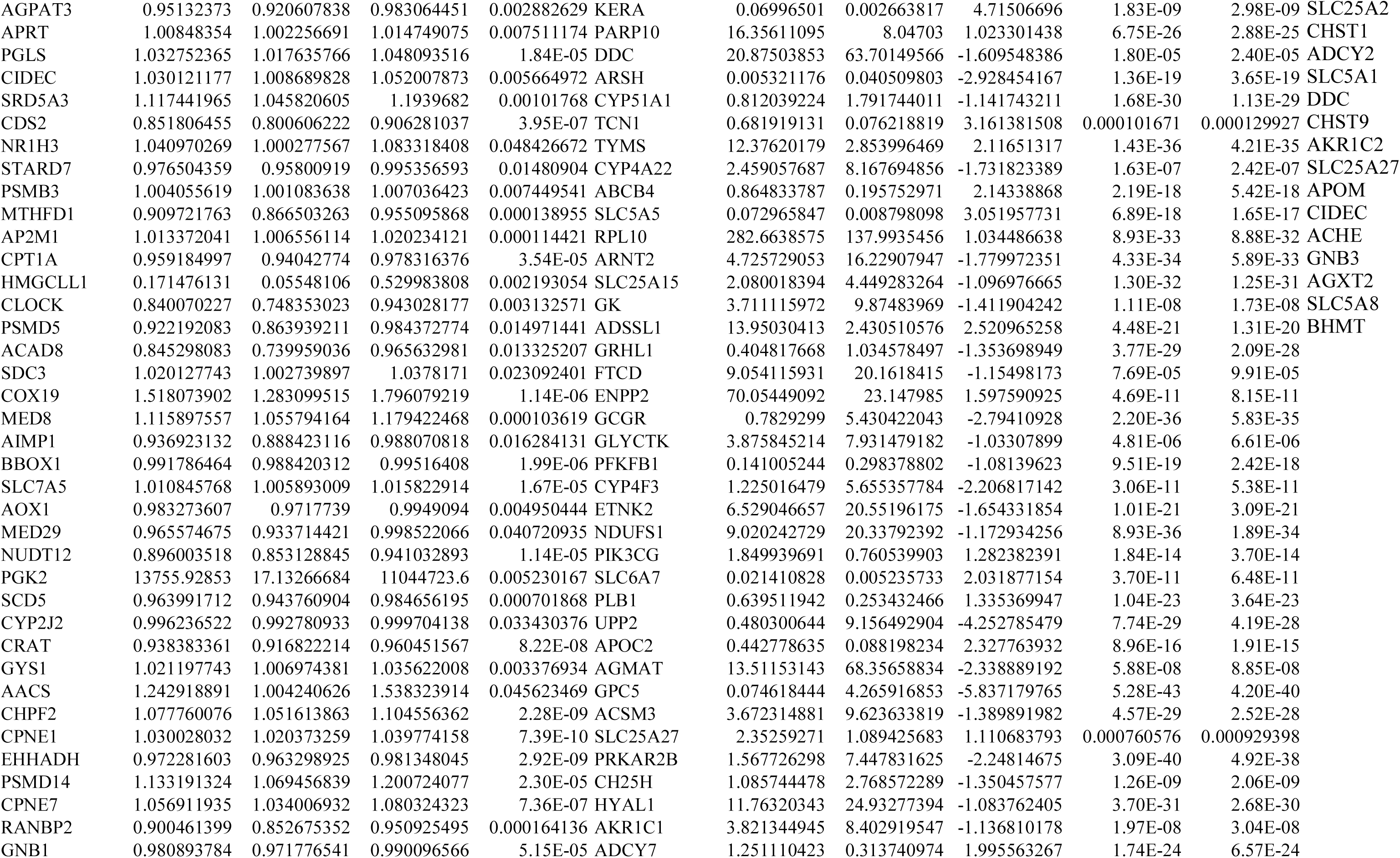
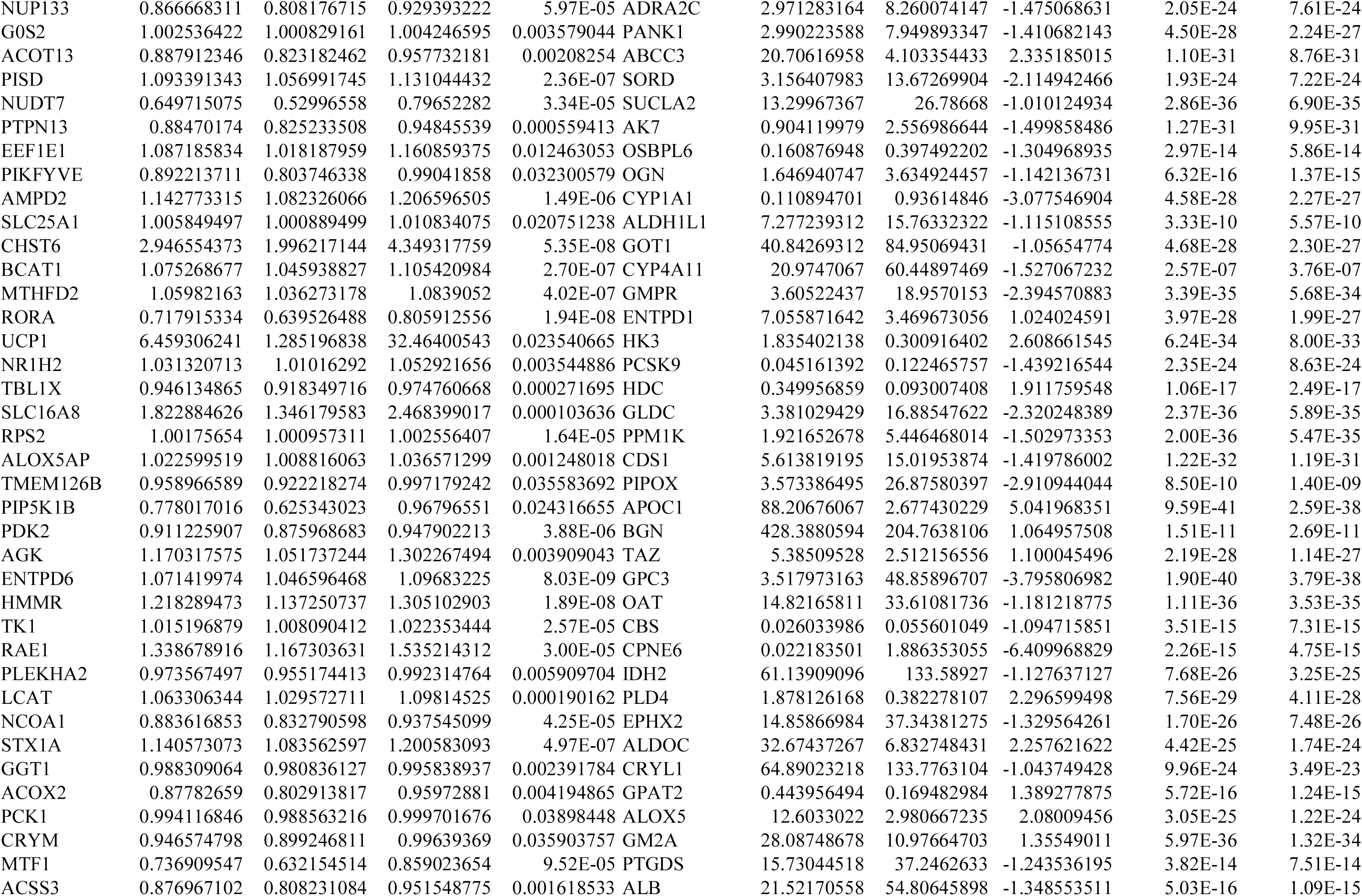
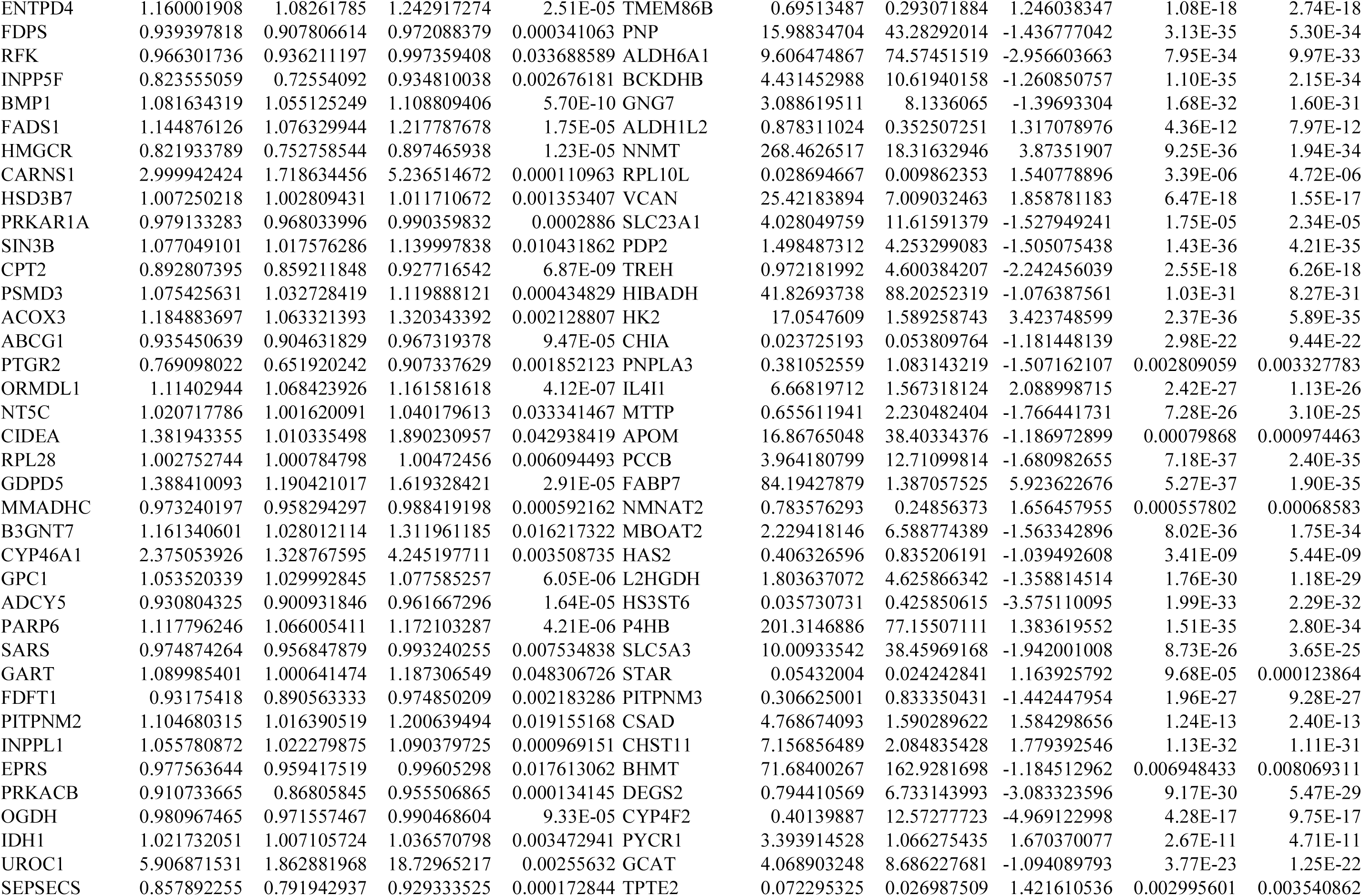
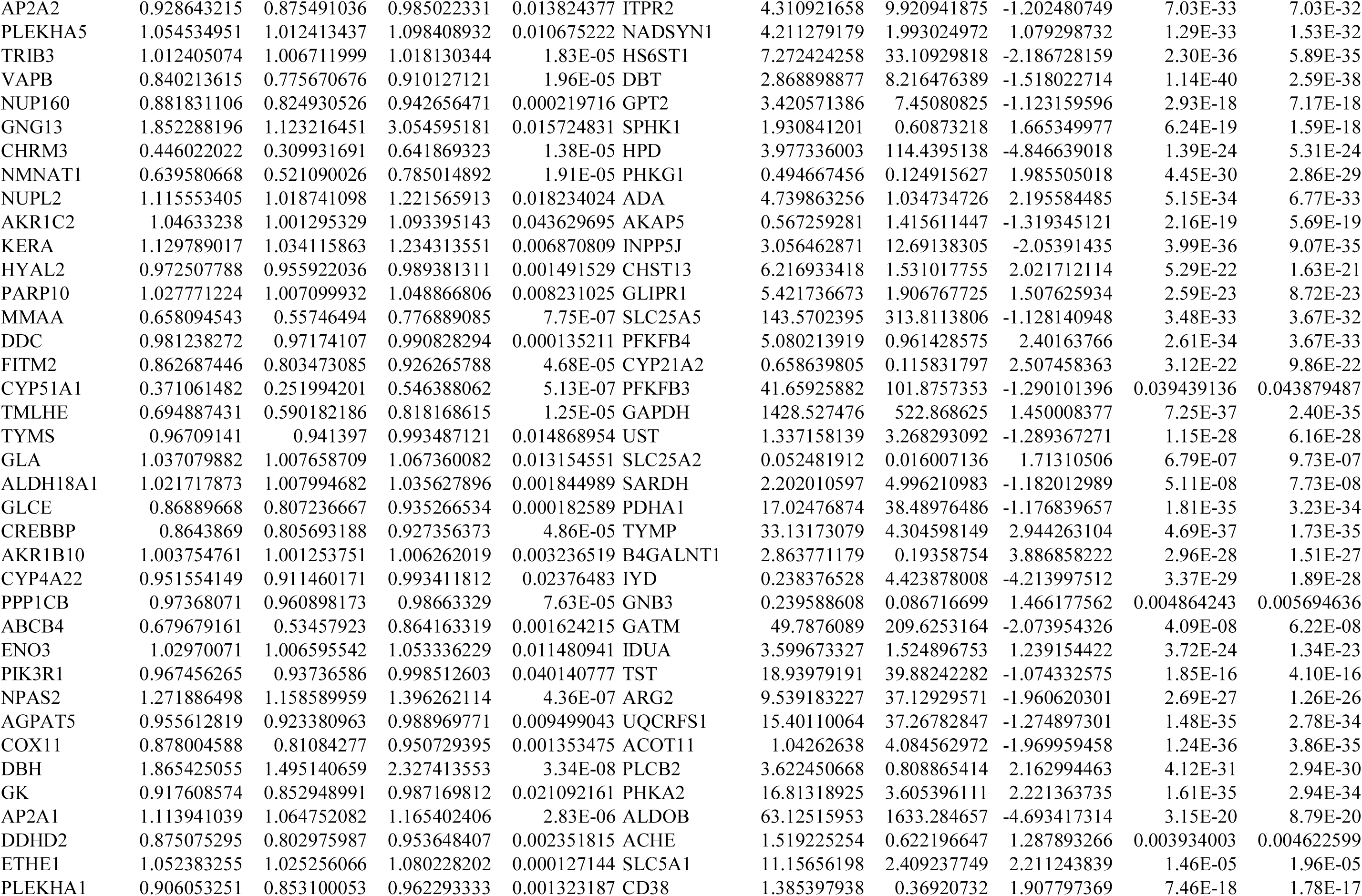
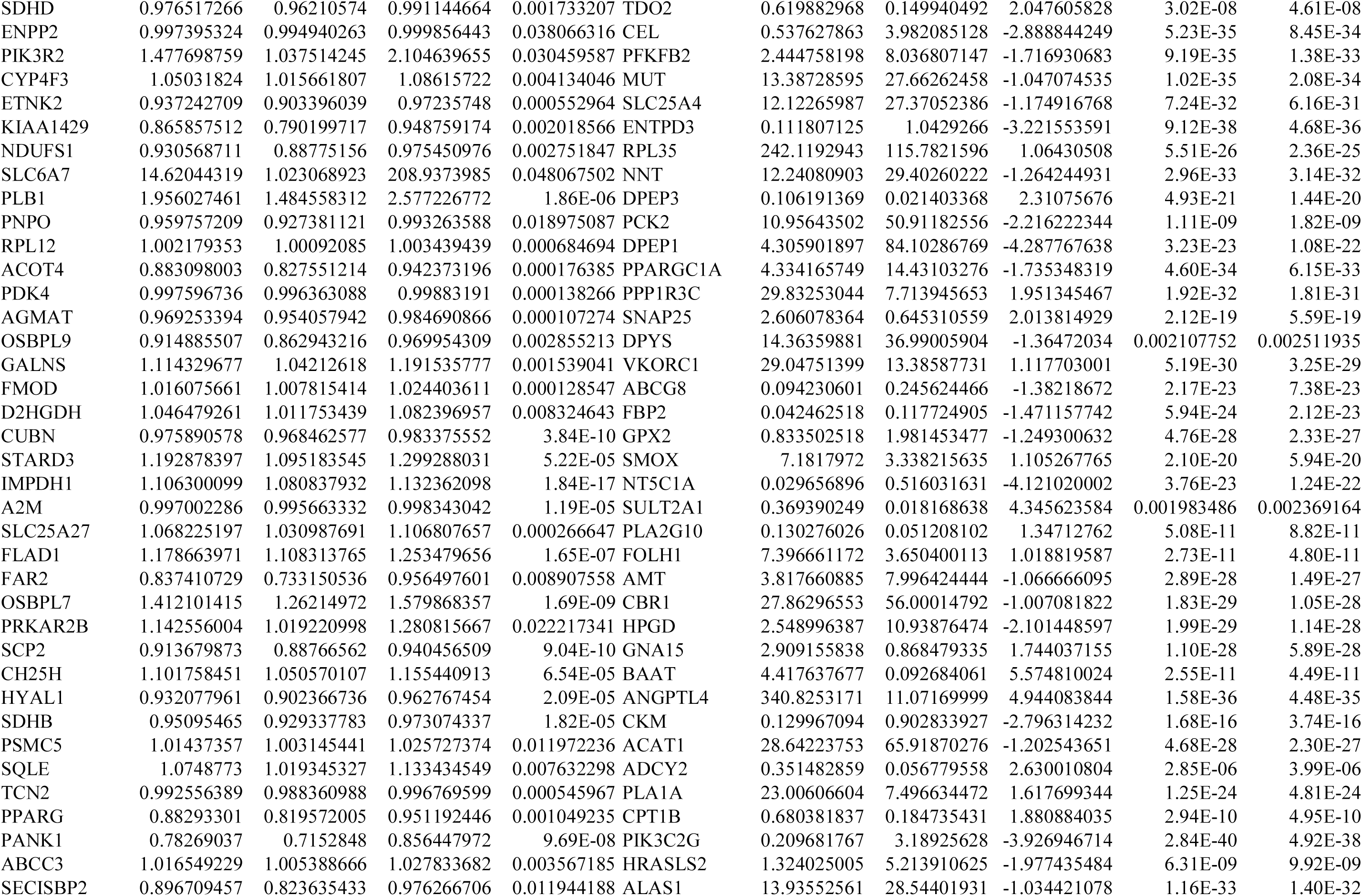
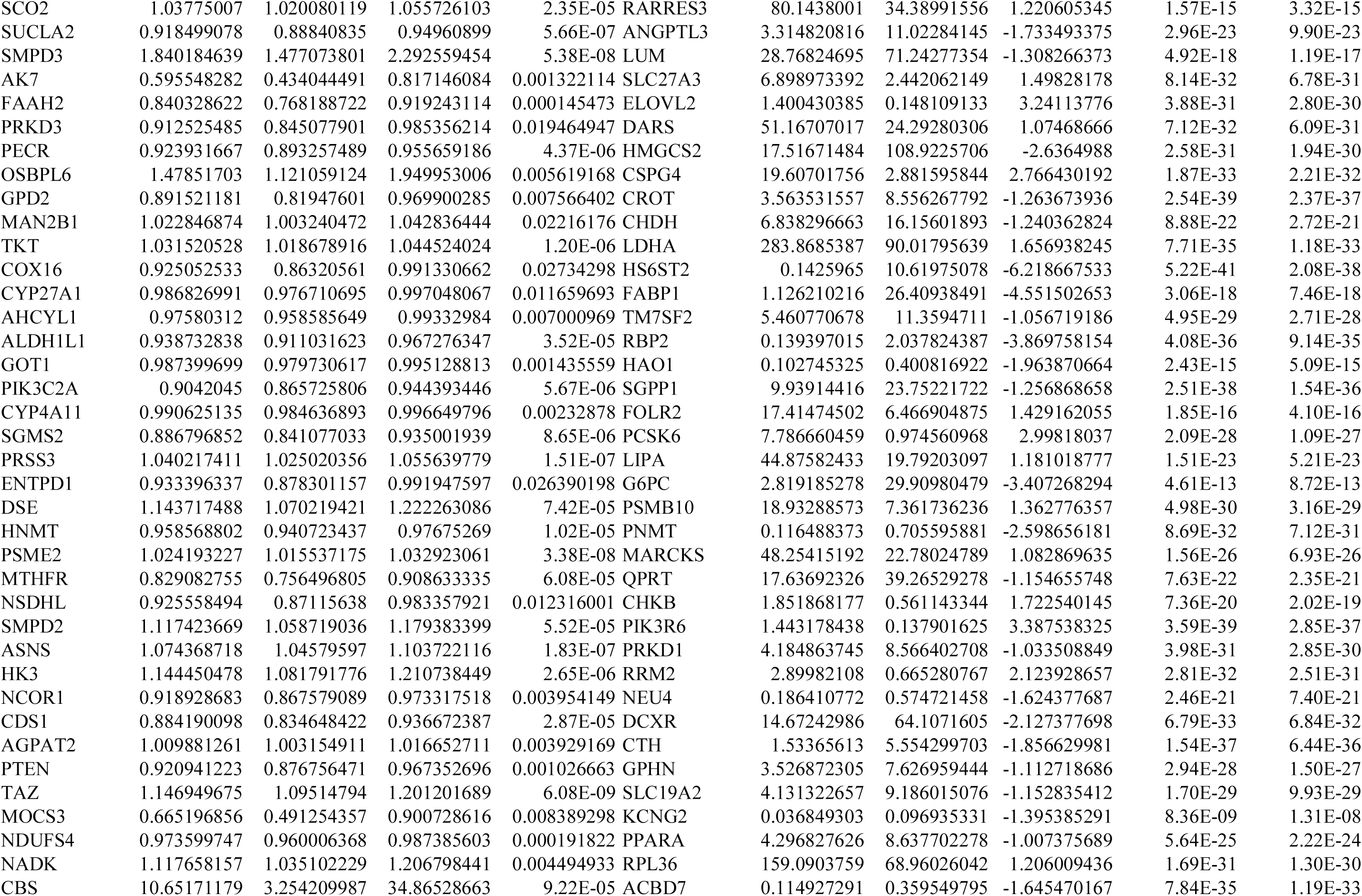
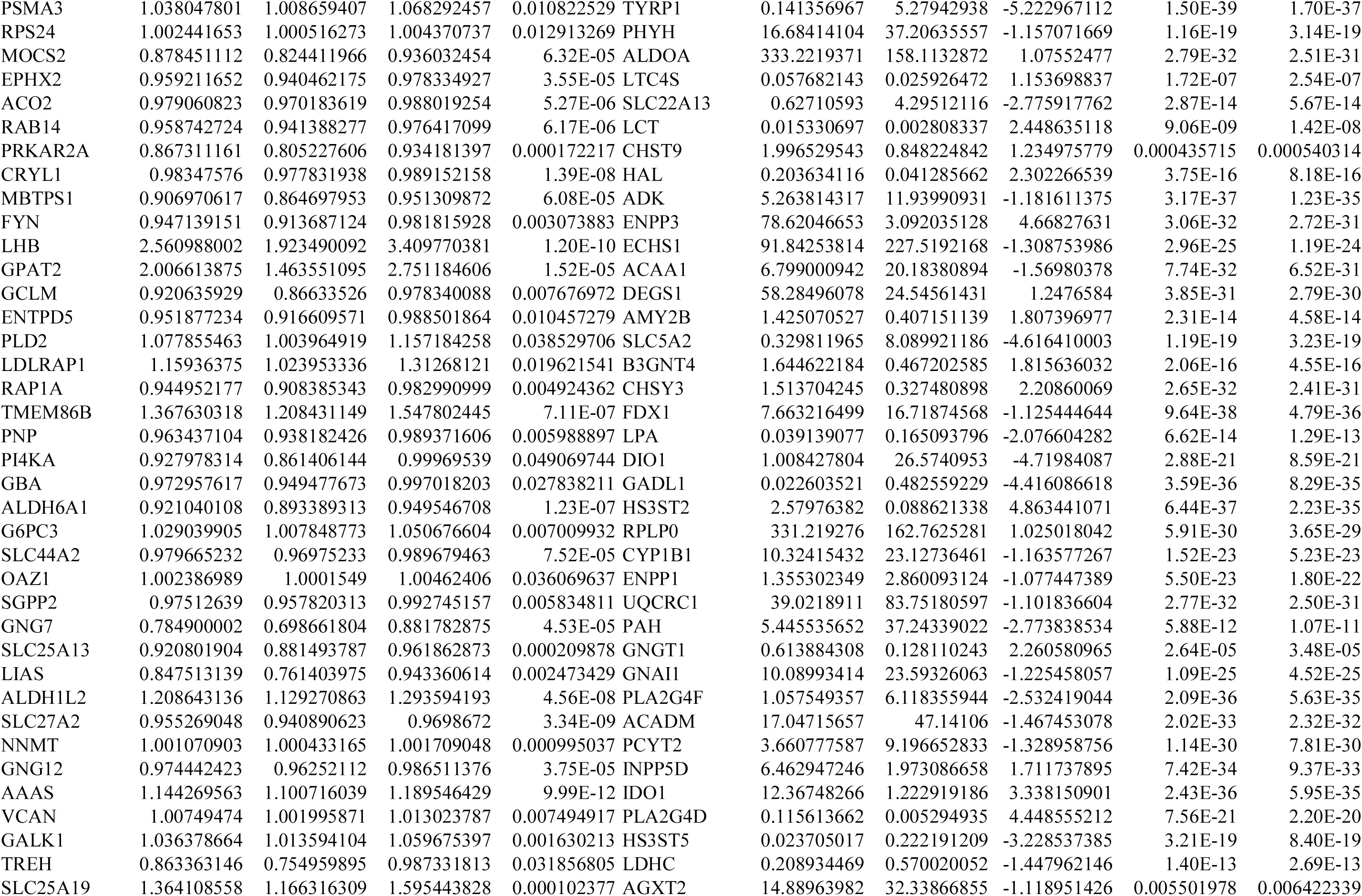
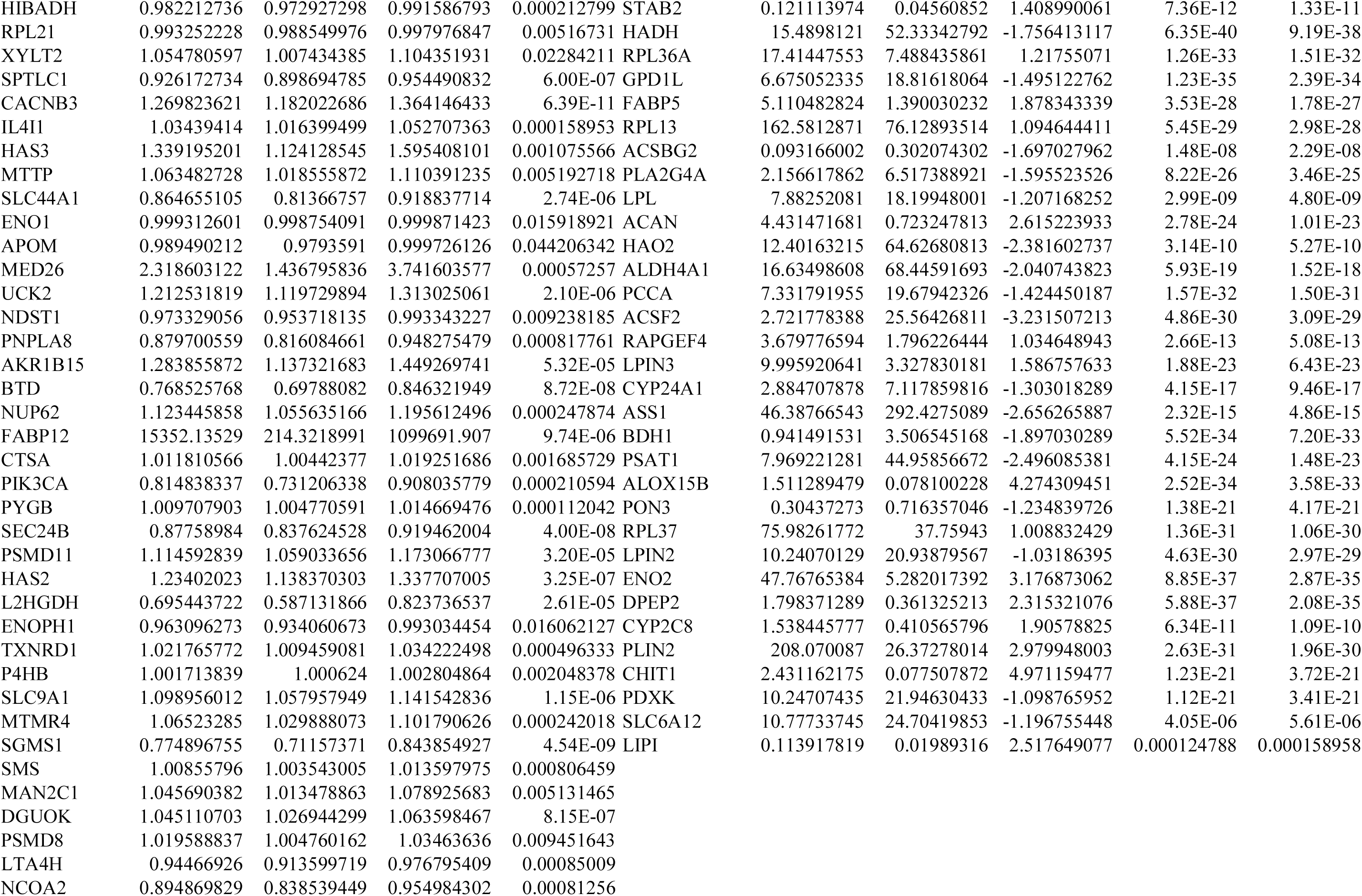
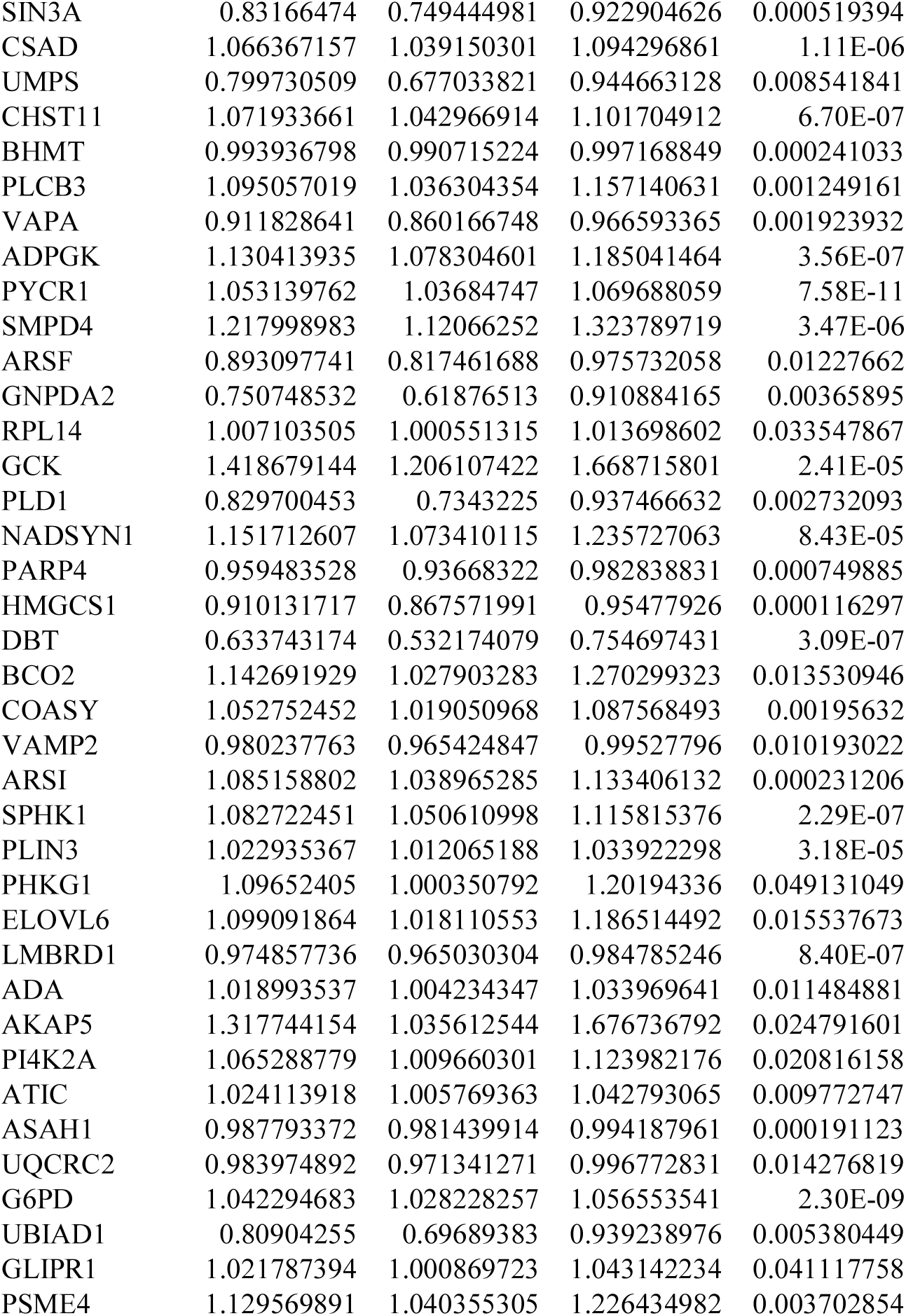
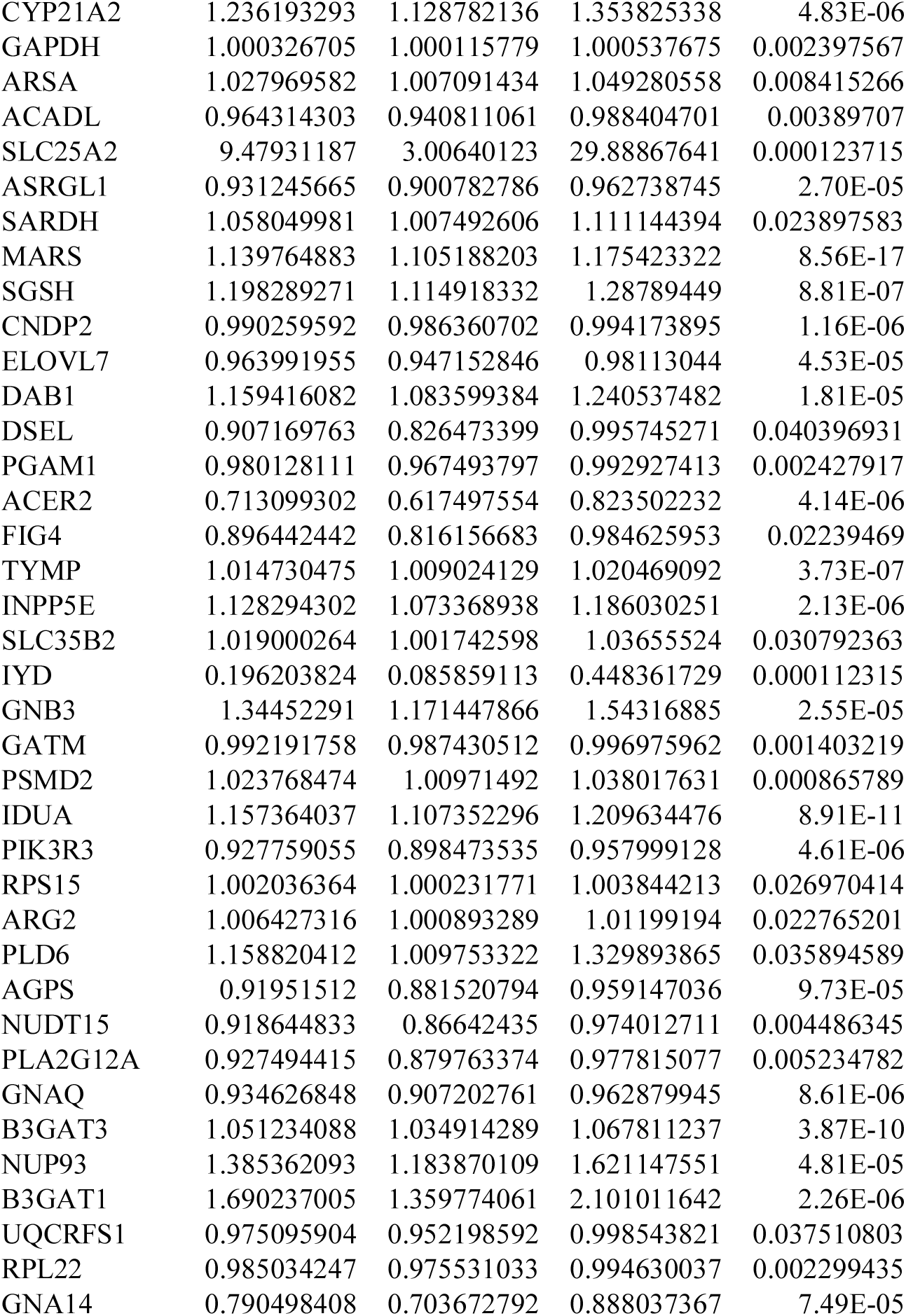
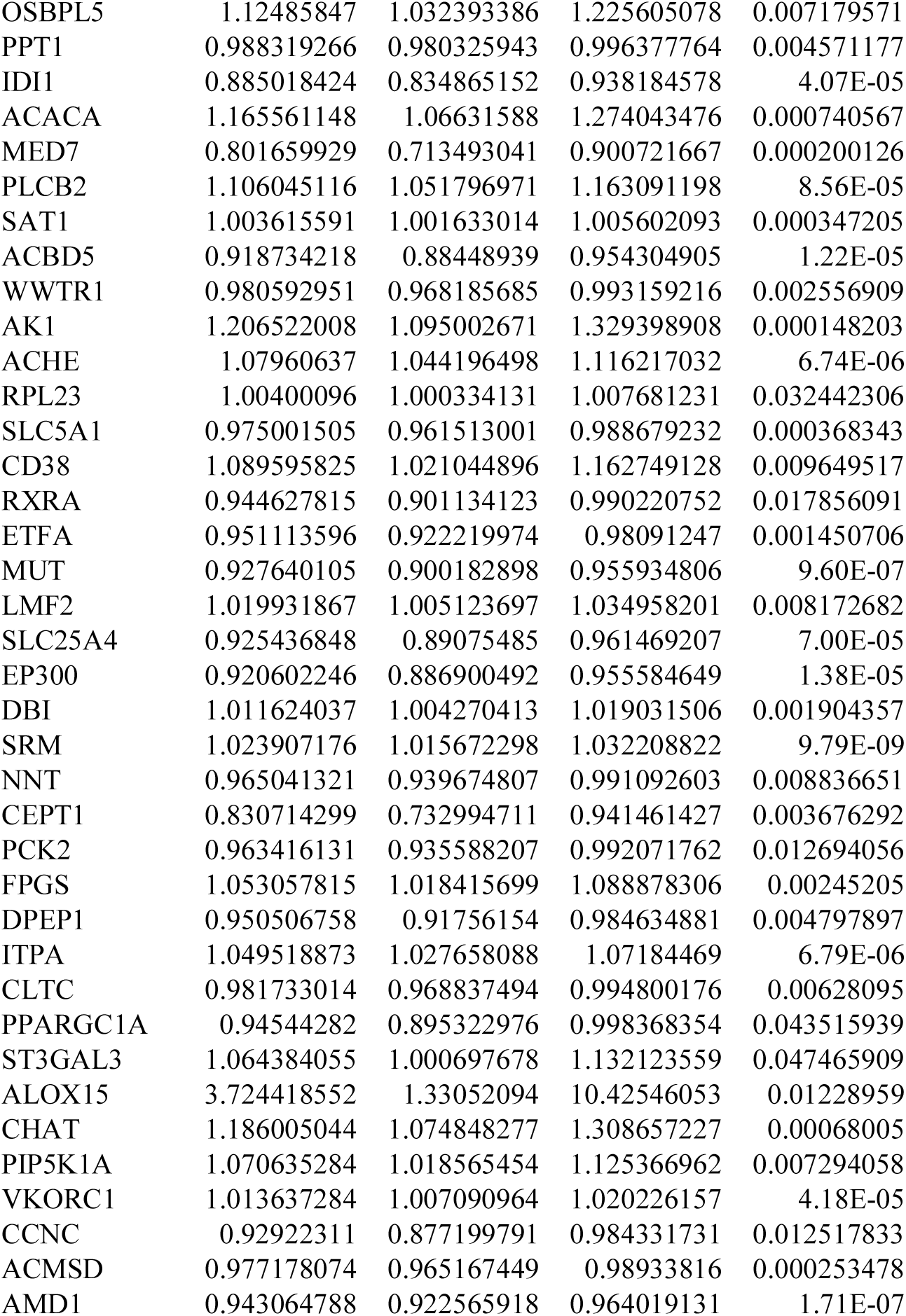
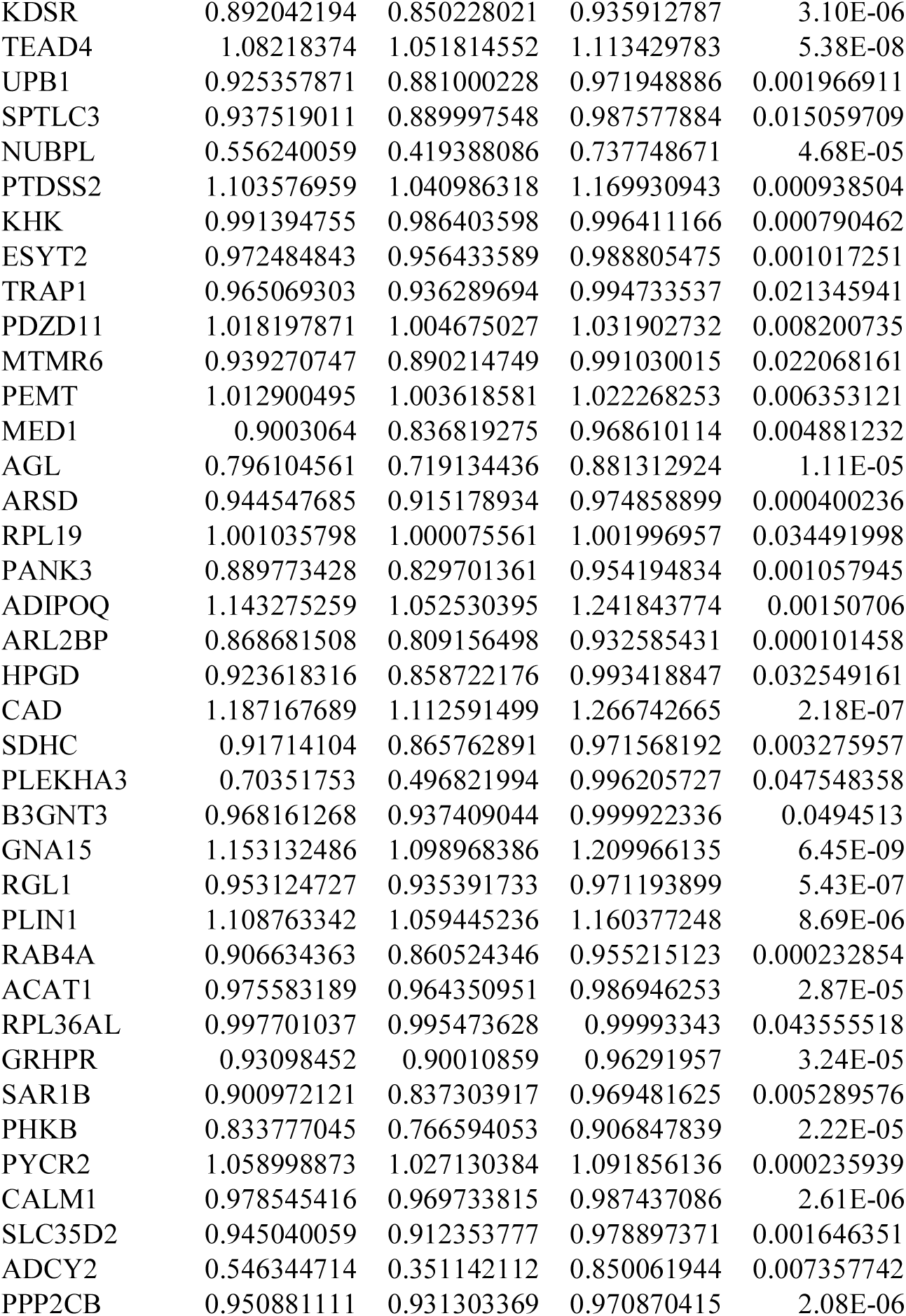
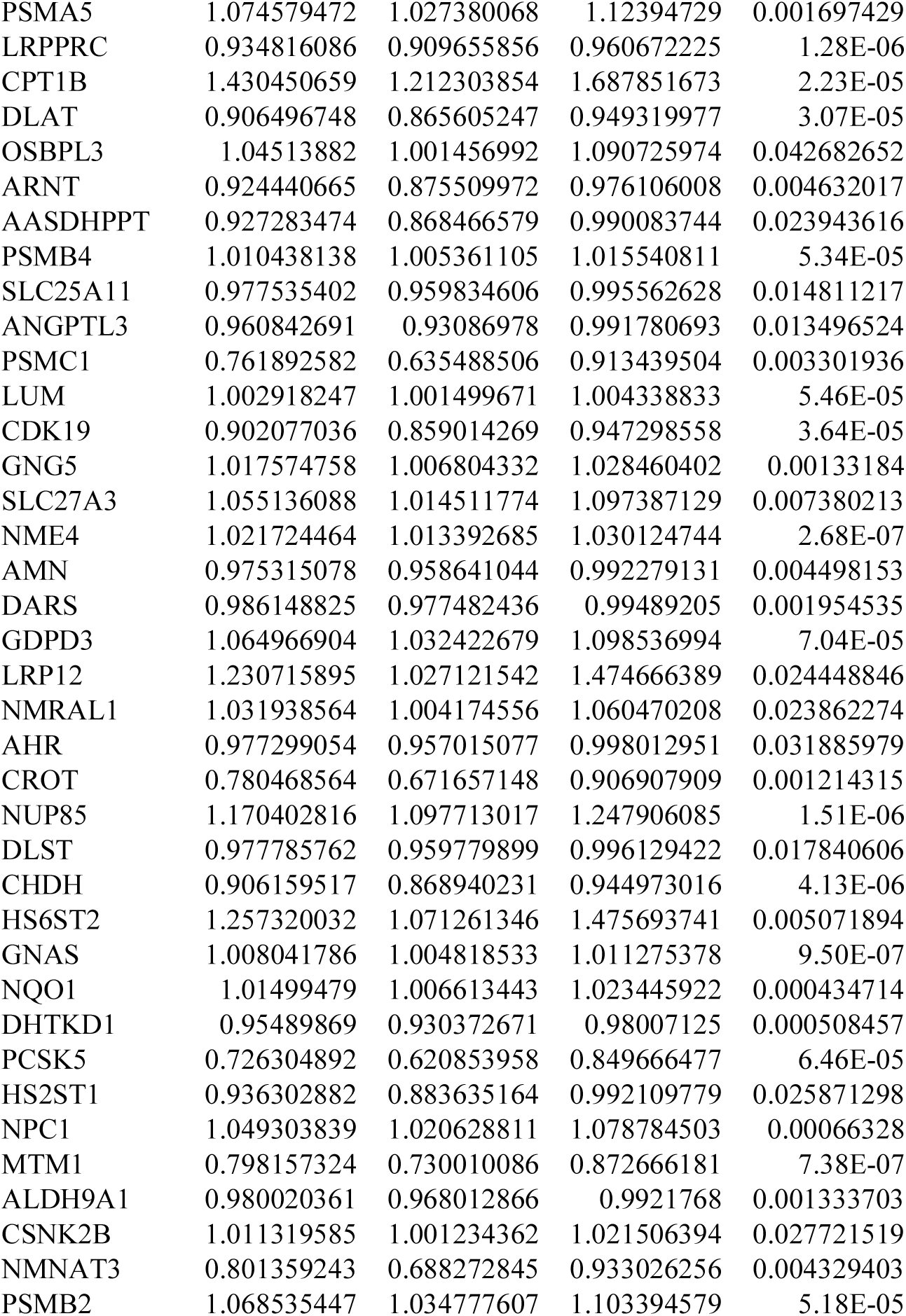
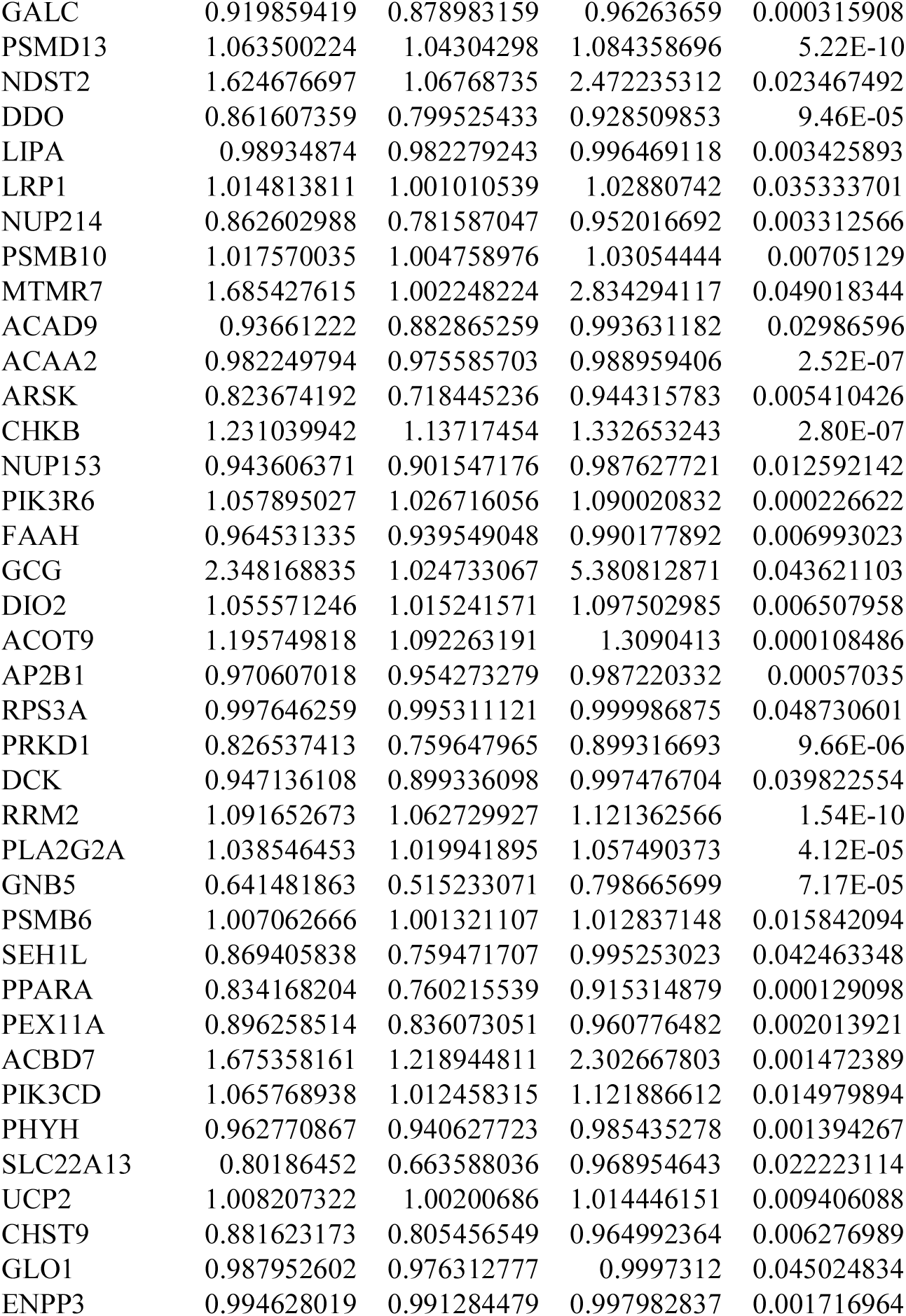
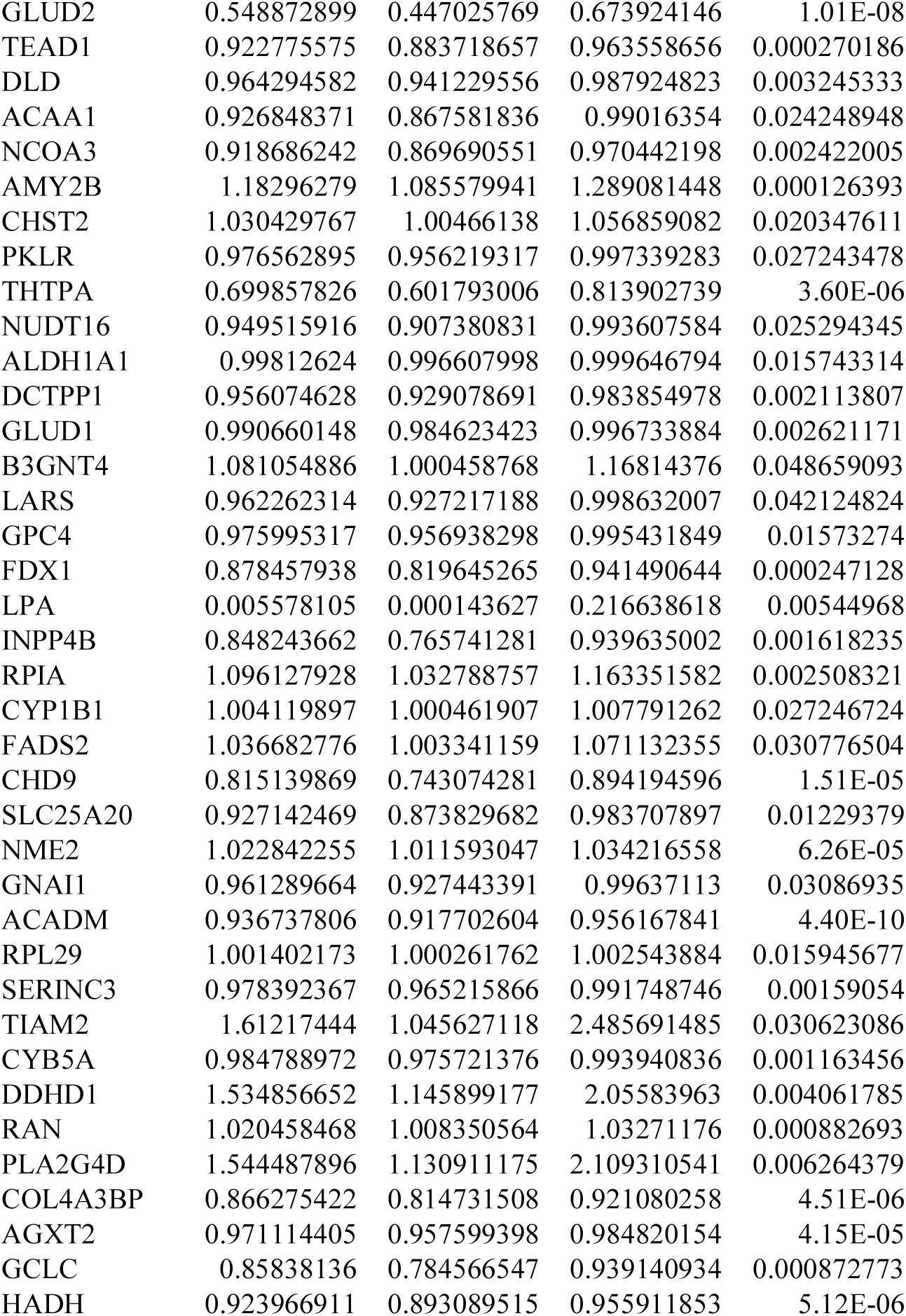
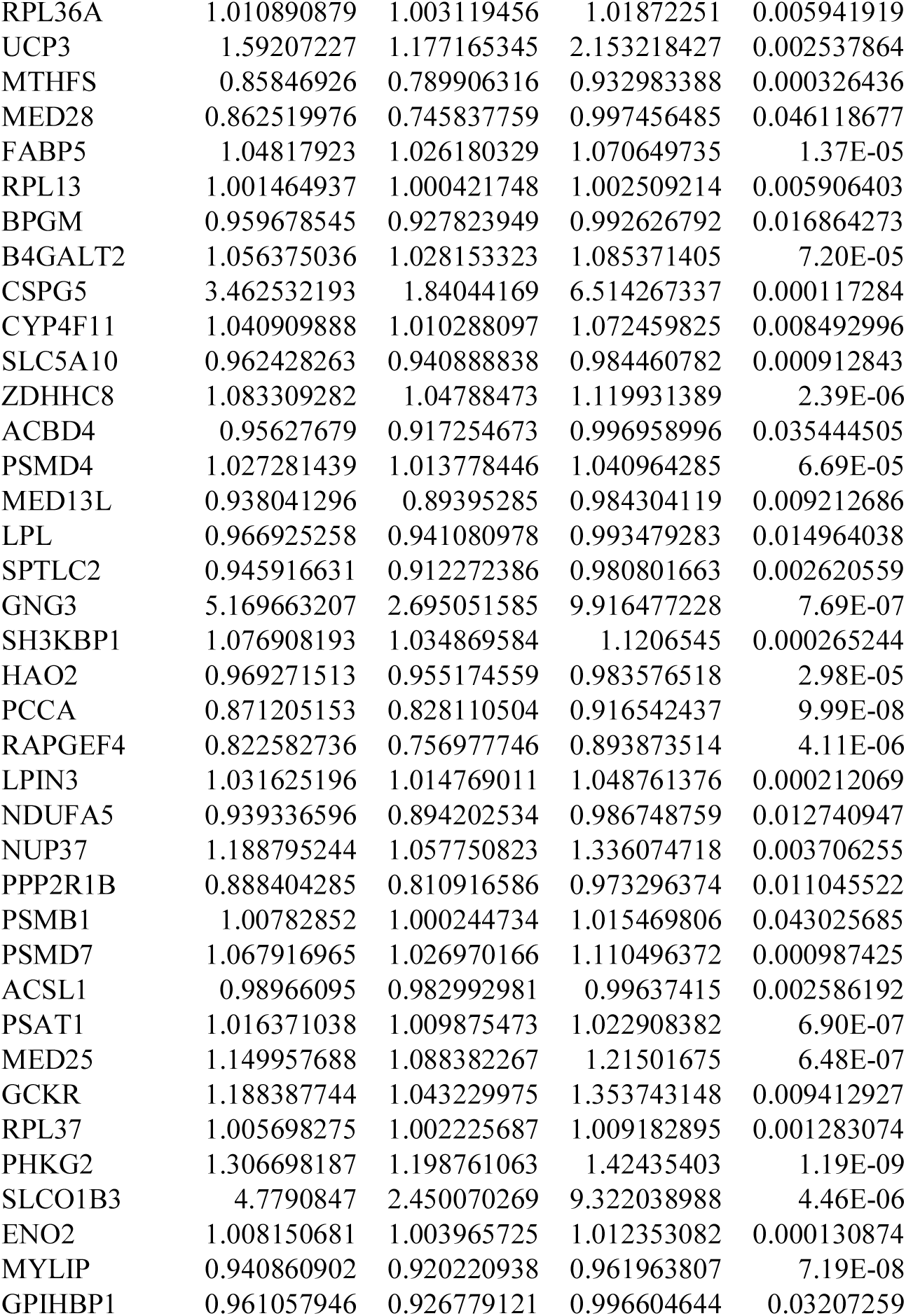
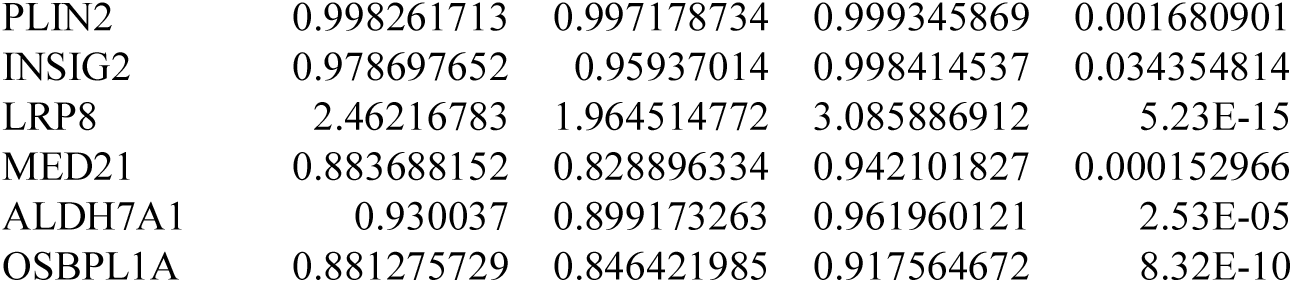

